# Holocene climate change promoted allopatric divergence and disjunct geographic distribution in a bee orchid species

**DOI:** 10.1101/2023.04.27.538532

**Authors:** Anaïs Gibert, Roselyne Buscail, Michel Baguette, Christelle Fraïsse, Camille Roux, Bertrand Schatz, Joris A. M. Bertrand

## Abstract

**Aim:** Species with disjunct geographic distributions provide natural opportunities to investigate incipient or recent allopatric divergence. The combination of both genetic and ecological data may be fruitful to decipher the causes of such pattern: i) actual vicariance, ii) successful colonization from one source range to a new range (dispersal, biological introduction) or iii) parallel convergent evolution.

**Location:** Southern France and Northern Spain.

**Taxon:** The bee orchid *Ophrys aveyronensis* (and its two recognized subspecies *O. a.* subsp. *aveyronensis* and *O. a.* subsp. *vitorica*) displays a disjunct geographic distribution with two subranges separated by 600 km on both sides of the Pyrenees Mountains.

**Methods:** Through a combination of population genomics and Ecological Niche Modelling (ENM), we investigated the causes of this intriguing biogeographic pattern.

**Results:** The population genomic data demonstrate that all the studied populations exhibit similar patterns of genetic diversity and dramatic decrease in effective size compared to the ancestral population. Significant genetic differentiation and reciprocal monophyly exist between populations of the two subranges of *O. aveyronensis*, despite a very recent divergence time as young as ca. 1500 generations ago. Moreover, (paleo-)ecological Niche Modelling analyses support that the disjunct geographic distribution of the *O. aveyronensis* is consistent with a range split of a broad ancestral range, contraction and distinct longitudinal and latitudinal shifts in response to climate warming during the Holocene.

**Main Conclusion:** The congruence of the results obtained from both population genomics and ENM approaches documents how very recent continental allopatric divergence initiated speciation in this system. *Ophrys aveyronensis* provides a promising opportunity to study the onset of reproductive isolation and parallel evolution following an initial stage of geographic separation in a group with high diversification rate.

## Introduction

Allopatric speciation (also referred as geographic speciation, or vicariant speciation) occurs when populations of a given ancestral species become reproductively isolated from each other because of geographic separation. Aside from being conceptually intuitive, allopatric speciation has been evidenced by theoretical models, laboratory experiments and empirical studies to the point that it is generally regarded as the “default mode” of speciation (see Coyne & Orr, 2004). In allopatric speciation, spatial and reproductive isolation are often assumed to originate from geologic-caused topographic changes (e.g. mountain formation) and/or climate-induced changes (e.g. sea level modifications in continental island systems) that form the geographic barrier. However, not only the fore-mentioned barriers but any kind of fragmentation of species distribution range, including those caused by human activities, may also initiate and be responsible for evolutionary divergence.

The putative complexity of the speciation process makes it difficult to elucidate its ecological and evolutionary causes, first because current spatial patterns of distribution only provide a snapshot from the speciation continuum (Nosil, Feder, Flaxman & Gompert, 2017; Stankowski & Ravinet, 2021). Allopatric divergence can be a spatio-temporally dynamic process as geographic and reproductive isolation is prone to be only transient. For example, climate change-induced oscillations during the Quaternary (Hewitt, 2000) have been shown to be an important driver of plant speciation and radiations, particularly in mountainous and arid environments (Kadereit & Abbott, 2021). In this context, populations may experience at least one phase of isolation and secondary contact which may alter our ability to properly infer the chronology of the divergence (see Ravinet et al., 2017) and separate the ‘historical- geographical’ from ‘ecogeographic’ causes of reproductive isolation (sensu Sobel, Chen, Watt & Schemske, 2010).

In this regard, taxa with disjunct geographic distribution may provide particularly insightful opportunities to study populations that became recently allopatric, *i.e.* with insufficient time to accumulate enough divergence to be considered as distinct species yet, as promising candidates to investigate incipient parallel evolution (see James et al., 2023). However, reviews of empirical cases of speciation tend to point out the relative scarcity of such candidates having started to diverge as recently as during the Holocene, especially in plants (see Kadereit & Abbott, 2021; de Queiroz et al., 2022). Aside, although such natural example may help to understand the consequences of physical barriers in early stages of evolutionary divergence, disjunct geographic distributions do not necessary result from the fragmentation of an ancestral range. Indeed, dispersal and successful colonization of a new range from the historical one may also lead to a similar biogeographic pattern.

The species of the genus *Ophrys* have long been known for their unusual pollination syndrome (called ‘sexual swindling’) that involves insect-like flowers mimicking their female pollinator to lure conspecific males and ensure pollination but are now also increasingly considered as a promising biological model to investigate adaptive radiations. The degree of plant-pollinator specialization (Joffard, Massol, Grenié, Montgellard & Schatz, 2019) required to ensure the success of the *Ophrys* pollination strategy was shown to be the driver of the impressive rate of diversification of the genus within the Mediterranean basin, and to a lesser extent in other parts of Western Europe (see Baguette, Bertrand, Stevens & Schatz, 2020, for a recent review). If *Ophrys* are notorious for their relatively high number of likely cases of speciation with gene flow, the number of island endemic taxa suggests that geographic isolation (induced by insularity) is also a key factor to form the large number of recent and emerging *Ophrys* species observable nowadays (see Bertrand, Baguette, Joffard & Schatz et al., 2021).

On continents, assessing the role of allopatry in *Ophrys* diversification is more challenging, in particular because clear evidences of taxa displaying subranges without being connected by gene flow are lacking. Nevertheless, one species, *Ophrys aveyronensis* (J.J. Wood) P. Delforge 1984, initially described as *Ophrys sphegodes* subsp. *aveyronensis* J.J. Wood,1983, displays an intriguing case of disjunct geographic distribution. This taxon was initially described as endemic to a very restricted area in the Grands Causses region (Southern karstic areas of the Massif Central, France). Observations of morphologically very similar populations were reported later from mid-elevations regions in Northern Spain (La Rioja, Burgos, Alava and Vizcaya provinces) and described as a distinct species (*Ophrys vitorica*, Kreutz, 2007). *Ophrys aveyronensis* is now rather considered as comprising two subspecies (*O. aveyronensis* subsp. *aveyronensis* and *Ophrys aveyronensis* subsp. *vitorica*) that share, at least a mutual insect pollinator species (*i.e.* the solitary bee *Andrena hattorfiana*, Paulus & Gack, 1999; Paulus, 2017; Benito Ayuso, 2019) but show subtle differences in morphometry and coloration (Gibert et al., 2022). This taxon is thus present in two rather restricted ranges separated by a gap of about 600 km (and Pyrenees Mountains), in the Grands Causses region (France) and in the central part of Northern Spain (Fig. 1).

**Figure 1.**
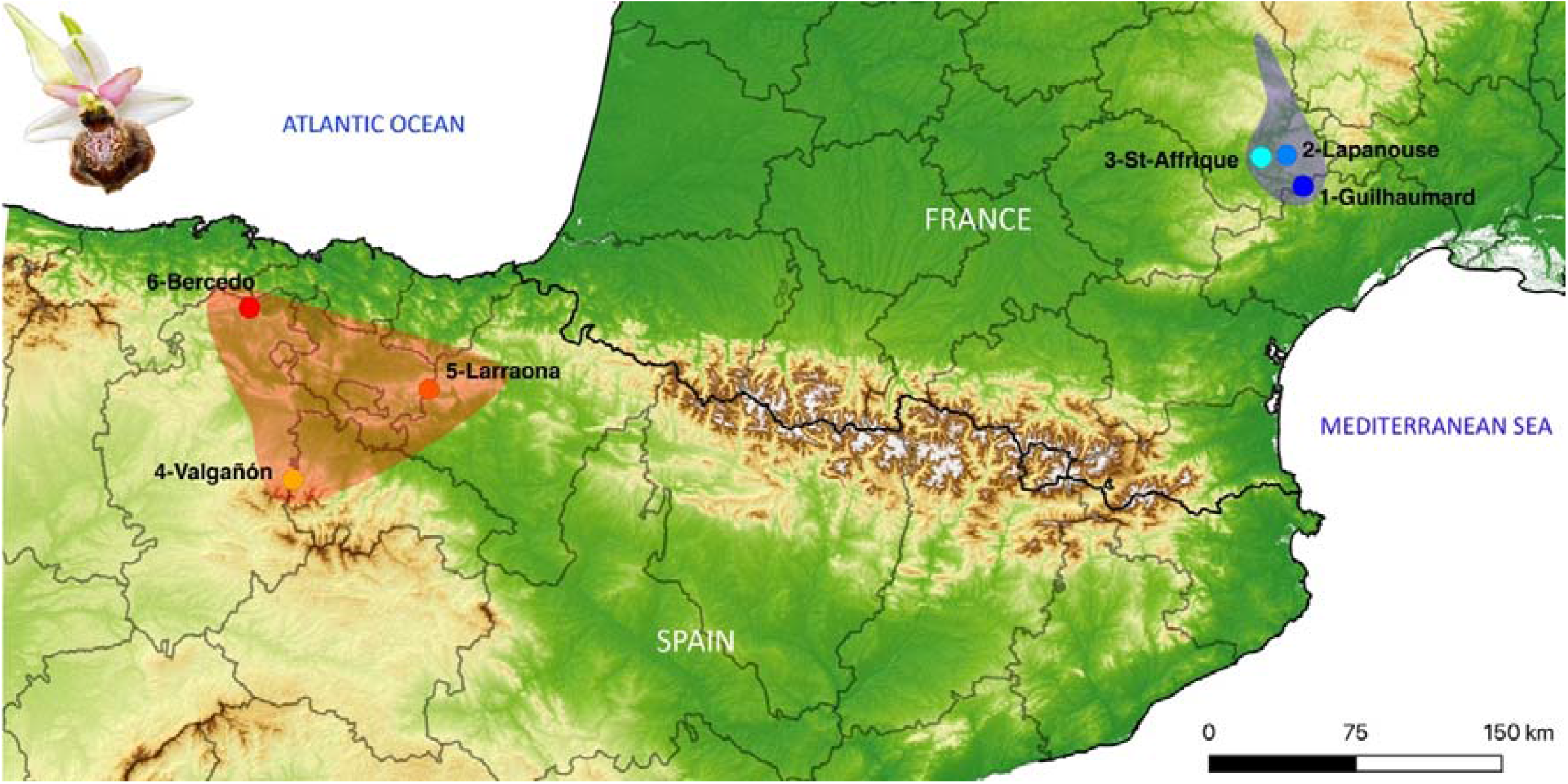
Map displaying the 6 sites sampled: 3 for *Ophrys aveyronensis* subsp. *aveyronensis*: 1-Guilhaumard, 2-Lapanouse-de-Cernon, 3-St-Affrique (in blue) and 3 for *O. a.* subsp. *vitorica*: 4- Valgañón, 5-Larraona, 6-Bercedo (in red). The approximate known geographic distribution is highlighted for both taxa with the same color code. The photograph (on the top left corner) illustrates the insect-mimicking flower of a typical *O. aveyronensis* with a marbled labellum.

The formation of the Pyrenees during the Eocene (from 45 Myrs ago) cannot explain by itself the split of the ancestral range of the species as all phylogenetic hypotheses published so far conclude that the *Ophrys* genus emerged no more than 5-6 Myrs ago and that the *Sphegodes* clade, to which *O. aveyronensis* belongs may have diversified over the last million year (Breitkopf et al., 2015, see also Inda, Pimentel & Chase et al., 2012). This does not rule out allopatry as factor to explain current disjunct geographic distribution but suggests that geographic separation would have occurred recently. According to this ‘vicariance hypothesis’, the current disjunct geographic distribution may correspond to two relict pockets of an ancestral range that got fragmented following a scenario according to which populations from Northern and Southern ranges diverge in allopatry. In this case, the degree of genetic differentiation between Northern and Southern populations is expected to be significant. Namely, genetic differentiation is expected to be relatively higher between Northern and Southern ranges than between populations within subranges. Alternatively, we also envisage a ‘biological introduction hypothesis’: a colonization from one historical range to the other. The notoriously small *Ophrys* seeds are likely to be wind-dispersed over hundreds of kilometers (*i.e.* thus being able to travel across most physical barriers including mountains and vast open water areas) and may have found in the newly colonized area, suitable conditions for their germination, development and reproduction. Although ‘orchid dusty seed rains’ may allow to generate founder populations with non-negligible genetic diversity, such newly established populations typically consist of a subset of the genetic pool of the source population (*i.e.* a relatively little divergent population with lower effective population size and genetic diversity). This could lead to a phylogenetic pattern where the populations of the founded taxon are nested within those of the source taxon, rendering the later paraphyletic with respect to the former.

In this study, we used a population genomic approach (*i.e.* through a double digest RAD-seq like protocol called ‘nGBS’) to first investigate patterns of genetic diversity and differentiation among *Ophrys aveyronensis* populations from both part of its disjunct geographic range. We then couple demographic inference analyses and Ecological Niche Modelling approaches to test whether the current disjunct geographic distribution of this species is rather consistent with either vicariance or recent dispersal. Specifically, we aimed at determining how recent the divergence between Northern Spain and Southern France populations of *O. aveyronensis* may be and whether or not this disjunct geographic distribution may have originated with a colonization event from an ancestral range to a new one.

## Materials and Methods

### Sampling and study sites

We used a minimally destructive sampling method to collect a total of 86 plant tissue samples from six localities that span the entire geographic range of *O. aveyronensis*: three in Southern France (Grands Causses region): 1-Guilhaumard, 2-Lapanouse-de-Cernon and 3- Saint-Affrique, and three in Northern Spain: 4-Valgañón, 5-Larraona and 6-Bercedo, in June 2019, Fig. 1). All the specimens were photographed and phenotyped at a suite of morphological traits (see Gibert et al., 2022). At each sampling site, we also collected 6 to 10, 0.6 mm diameter leaf punches from 13 to 15 individuals which were stored in 100% ethanol until DNA extraction (see Table 1 and Supplementary Table S1). As *Ophrys aveyronensis* is a globally rare and nationally protected plant species in France, sampling was done under permit ‘Arrêté préfectoral n°2019-s-16’ issued by the ‘Direction Régionale de l’Environnement de l’Aménagement et du Logement (DREAL)’ from the ‘Région Occitanie’, on 07-May-2019). *Ophrys aveyronensis* is not legally protected in Spain.

**Table 1.**
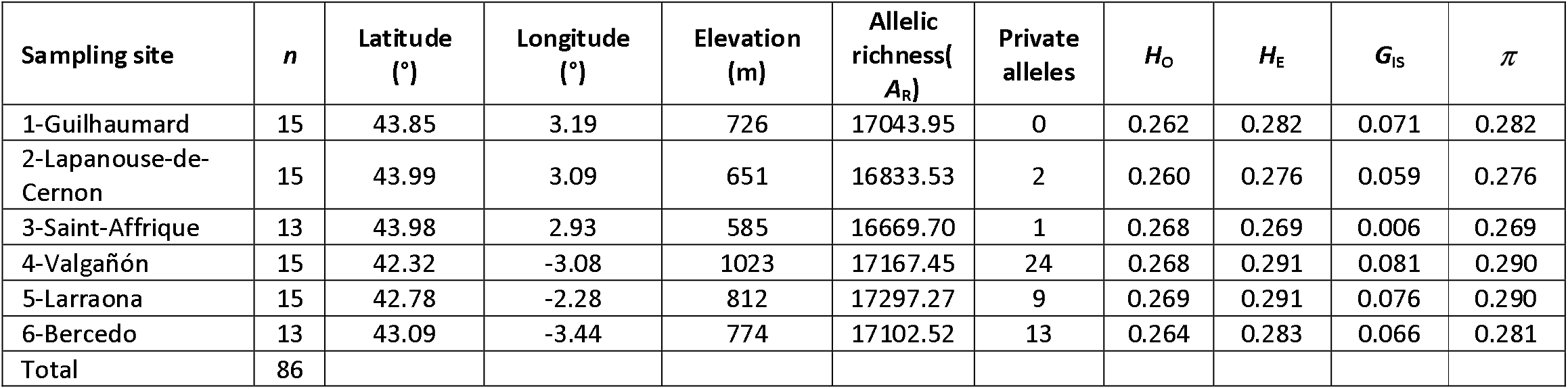
Sampling site, sample size (*n*), geographic coordinates and elevation, population summary statistics: allelic richness per sampling site (*A*_R_), number of private alleles, expected and observed heterozygosity (*H*_E_ and *H*_O_), inbreeding coefficient (*G*_IS_) and nucleotide diversity (*π*)

| Sampling site | $n$ | Latitude (°) | Longitude (°) | Elevation (m) | Allelic richness($A_R$) | Private alleles | $H_O$ | $H_E$ | $G_{IS}$ | $\pi$ |
| --- | --- | --- | --- | --- | --- | --- | --- | --- | --- | --- |
| 1-Guilhaumard | 15 | 43.85 | 3.19 | 726 | 17043.95 | 0 | 0.262 | 0.282 | 0.071 | 0.282 |
| 2-Lapanouse-de-Cernon | 15 | 43.99 | 3.09 | 651 | 16833.53 | 2 | 0.260 | 0.276 | 0.059 | 0.276 |
| 3-Saint-Affrique | 13 | 43.98 | 2.93 | 585 | 16669.70 | 1 | 0.268 | 0.269 | 0.006 | 0.269 |
| 4-Valgañón | 15 | 42.32 | -3.08 | 1023 | 17167.45 | 24 | 0.268 | 0.291 | 0.081 | 0.290 |
| 5-Larraona | 15 | 42.78 | -2.28 | 812 | 17297.27 | 9 | 0.269 | 0.291 | 0.076 | 0.290 |
| 6-Bercedo | 13 | 43.09 | -3.44 | 774 | 17102.52 | 13 | 0.264 | 0.283 | 0.066 | 0.281 |
| Total | 86 |  |  |  |  |  |  |  |  |  |

### DNA extraction, library preparation and genotyping

Genomic DNA extraction as well as library preparation and genotyping were subcontracted to LGC Genomics GmbH (Berlin, Germany). A fractional genome sequencing strategy called normalized Genotyping-by-Sequencing (nGBS) was used to subsample the *Ophrys aveyronensis* genome. Most *Ophrys* species are diploid: 2n = 36 (Turco, Albano, Medagli, Wagensommer & D’Emerico, 2023) and *Ophrys aveyronensis*, for which we estimated a relatively large size of ∼5-6 Gbp (1C-value = 5.44 to 6.02 picograms, *n* = 3 for *O. a.* subsp. *aveyronensis* and 1C-value = 5.88 pg, *n* = 1, for *O. a.* subsp. *vitorica*) based on flow cytometry techniques (Bertrand, unpublished data) conforms to previous knowledge. The nGBS protocol makes use of a combination of two restriction enzymes (PstI and ApeKI, in our case) to produce a reproducible set of fragments across samples/individuals. It also comprises a normalization step whose aim is to avoid further sequencing of highly repetitive regions. Illumina technology was then used to aim at obtaining a minimum of 1.5 million reads per sample (with 2 x 150 bp read length) for each of the 86 individually barcoded libraries.

### Population genomic dataset processing

We used different scripts included in *Stack*s v.2.60 (Catchen Amores, Creskp & Postlewait, 2011; Catchen, Hohenlohe, Bassham, Amores & Cresko, 2013) to build loci from Illumina reads, *de novo* (*i.e.* without aligning reads to a reference genome). We first used *process_radtags* to demultiplex and clean reads, then *denovo_map.pl* to build loci within individuals, create a catalog and match all samples against it and finally *populations* to further filter the SNPs obtained at the population level and compute basic population genetic statistics. Before running the pipeline on the complete dataset, we first optimized several key parameters: *-m* (the minimum number of identical raw reads required to form a putative allele), -*M* (the number of mismatches allowed between alleles to form a locus) and *-n* (the number of mismatches allowed between loci during construction of the catalog) by running it on a subset of 12 individuals representative of the whole dataset (*i.e.* geographic origin and coverage). As recommended by several authors, we varied *M* and *n* from 1 to 9 (fixing *M* = *n*) while keeping *m* = 3 (see Paris, Stevens, & Catchen, 2017). The combination -*m* 3, -*M* 5 and -*n* 5 was found to be the most suitable to maximize the number of SNPs, assembled and polymorphic loci in our case and was used to run *Stacks* on the whole data set (see Supplementary Table S2). Then, we ran the *populations* script, from which only SNPs that were genotyped in at least 80% of the individuals per population (-*r* 80) and found in all six sampling sites were kept (-*p* 6), while SNPs exhibiting estimates of observed heterozygosity greater than 70% (--*max-obs-het* 0.7) were filtered out to reduce the risk of including remaining paralogs. We also discarded sites whose minor allele frequency was lower than 5% (--*min-maf* 0.05). The genotype matrix of the remaining 9301 SNPs was then exported in several formats for downstream analyses.

The following population genomic analyses aimed to i) infer patterns of genetic diversity and divergence between subspecies/countries and sampling sites and ii) infer the demographic scenario (and associated parameters) likely to best explain the current biogeographic pattern observed in the *O*. *aveyronensis*.

### Genetic diversity, population differentiation and evolutionary divergence

To get an overview of the overall genetic diversity and differentiation among individuals and populations, we first performed a Principal Component Analysis (PCA) based on a matrix of 86 individuals (as rows) and 9301 SNPs (as columns) coded as a *genlight* object with the R-package *adegenet* (Jombart, 2008; Jombart & Ahmed, 2011). The number of private alleles and the average nucleotide diversity (*π*) were directly available from the output of the *populations* script from *Stacks*. We used Genodive v.3.04 (Meirmans, 2020) to compute expected and observed heterozygosity (*H*_E_ and *H*_O_, respectively) as well as to evaluate the deviation from panmixia by computing *G*_IS_. Allelic richness per sampling site (*A*_R_) was computed with the R-package *hierfstat* (Weir & Goudet, 2017). Overall and pairwise genetic differentiation was also assessed based on *G*-statistics (*G*_ST_, *G*’_ST_, *G*’’_ST_) as well as Jost’s *D* also implemented in Genodive. Genodive was also used to carry out a hierarchical Analysis of Molecular Variance (AMOVA) to assess proportions of genetic variance (i) among groups (here, subspecies/countries) (*F*_CT_) and (ii) among populations within each subspecies (*F*_SC_). The statistical significance of the obtained values was estimated based on 10000 permutations.

To further investigate population structure and characterize putative migration/admixture event, we used sNMF (Frichot, Mathieu, Trouillon, Bouchard & François, 2014) as implemented in the R-package *LEA* (Frichot & François, 2015) to estimate individual ancestry coefficients based on sparse non-negative matrix factorization algorithms. The number of ancestral populations (genetic clusters) was varied from *K* = 1 to 10, and analyses were run with 10 replicates at each value of *K*. We followed the cross-entropy criterion to determine the optimal number of clusters and the ancestry proportions matrices (*Q*-matrices) obtained were plotted with the R-package *pophelper* (Francis, 2017).

Finally, we performed Maximum-Likelihood phylogenetic tree reconstruction with IQ-Tree v.2.2.6 (Minh *et al*., 2020) based on the concatenated SNP matrix with 1000 replicates (-B 1000) of Ultrafast Bootstrap Approximation (UFBoot) to assess nodes support. The substitution model that best fitted the data was selected with *ModelFinder Plus* (Kalyaanamoorthy, Minh, Wong, Von Haeseler & Jermiin, 2017) and used Ascertainment bias correction (-MFP+ASC) as recommended for SNP data.

### Demographic and evolutionary history reconstruction

We reconstructed the past demographic history of each of the six populations, separately, using STAIRWAYPLOT2 (Liu & Fu, 2015; 2020). This program relies on the Site Frequency Spectrum (SFS) to infer the temporal dynamics of effective population size (*N*_e_) and has the advantage of not requiring whole-genome sequence data or reference genome information. First, we used the *easySFS.py* script (https://github.com/isacovercast/easySFS) to compute the folded SFS of each population. To maximize the number of segregating sites, the populations were downsampled to 24, 24, 22, 24, 24 and 22 individuals for 1-Guilhaumard, 2-Lapanouse, 3-St-Affrique, 4-Valgañón, 5-Larraona and 6-Bercedo, respectively. We then ran STAIRWAYPLOT2 that fits a flexible multi-epoch demographic model that estimates *N*_e_ at different time periods using the SFS to infer estimated *N*_e_ fluctuation coinciding with coalescent events. Here, we assumed a mutation rate of 7 x 10^-9^ per site per generation for angiosperms autosomal markers (following Krasovec, Chester, Ridout & Filatov, 2018) a mean generation time of 5 years for *Ophrys*, and performed 200 bootstrap replicates to estimate 95% confidence intervals.

We then used the Approximate Bayesian Computation (ABC) framework implemented in DILS (Fraïsse et al., 2021) to infer the best demo-genomic scenario and associated parameters of the evolutionary history between *O. a.* subsp. *aveyronensis* and *O. a.* subsp. *vitorica*. We considered two types of datasets i) a pooled dataset grouping the three populations *O. a.* subsp. *aveyronensis* together and the 3 populations of *O. a.* subsp. *vitorica* together and ii) paired datasets consisting of all possible *O. a.* subsp. *aveyronensis* and *O. a.* subsp. *vitorica* population pairs (3^2^ = 9 pairwise comparisons). The two-population model of DILS compares different scenarios in a hierarchical manner. First, current isolation models, *i.e.* Strict Isolation (SI) versus Ancient Migration (AM) are compared to ongoing migration models, *i.e.* Isolation-with-Migration (IM) versus Secondary Contact (SC). Then, for the best demographic model, DILS compares genomic models to test for linked selection by assuming variation in effective population size (*N*_e_) across sites (*i.e.* hetero versus homo *N*e models are compared). For migration models (IM and SC), DILS also tests for selection against migrants by assuming variation in migration rates *N.m* across sites (*i.e.* hetero versus homo *N.m* models are compared). Depending on the best selected model, DILS provides estimates and confident intervals for parameters such as the time of split (*T*_split_), the migration rate (*N.m*), current and ancestral effective population sizes (*N*e_current_ and *N*e_past_) and whenever relevant, the time of ancient migration (*T*_AM_) or the time of secondary contact (*T*_SC_). We formatted the input file with a custom Perl script and used the online facility (https://www.france-bioinformatique.fr/cluster-ifb-core/) to run computations with the following settings and priors: *population_growth*: constant, *modeBarrier*: bimodal (*i.e.* there is a class of species barrier loci with *N.m* = 0 and a class of loci migrating at the background rate *N.m*), *max_N_tolerated:* 0.1*, LMin* = 30, *nMin* = 10 (20 for the pooled dataset), *mu* = 7.10^-9^, *rho_over_theta*: 0.1, *N*_min = 100, *N_max*: 200000, *Tsplit_min*: 100, *Tsplit_max*: 25000, *M_min*: 0.4 and *M_max*: 4. To quantify the fit of the best model to the data we conducted a goodness-of-fit test using 2000 simulations performed under the best model based on the estimated parameter values.

### Past and current Species Distribution Modelling

We used an Ecological Niche Modelling (or Environmental Niche Modelling, ENM or Species Distribution Modelling, SDM) framework similar to the one we followed and described in detail in Salvado et al. (2022) to infer the spatio-temporal dynamics of suitable habitat for *O. aveyronensis* since the Last Glacial Maximum (LGM, 21 kyrs BP). We then compared the results with those obtained from population genomic analyses. To model and spatially project the current bioclimatic niche, we first downloaded 19 bioclimatic layers at a resolution of 2.5 arc-minutes (∼5 km^2^) available from WorldClim 2 (Fick & Hijmans, 2017). Current climate data correspond to time averaged variables over the period 1970-2000. To spatially project the inferred niche onto past bioclimatic conditions data, we then downloaded paleoclimate data either from WorldClim v1.4 (Hijmans, Cmaeron, Para, Jones & Jarvis, 2005) or from PaleoClim (www.paleoclim.org).

Paleoclimate data from WorldClim are available for both the LGM (21 kyrs ago) and the Mid-Holocene (about 6 kyrs ago), downscaled at a resolution of 2.5 arc-minutes for at least three commonly used General Circulation Models (GCMs) proposed by the National Center for Atmospheric Research (NCAR) though the Community Climate System Model (CCSM4), the Japan Agency for Marine-Earth Science and Technology, the Center for Climate System Research of the University of Tokyo, and the National Institute for Environmental Studies (MIROC-ESM) and the Max Plant Institute for Meteorology Earth System Model (MPI-ESM-P). Only CCSM data are available from PaleoClim but over different time periods over the Pleistocene (at a resolution of 2.5 arc-minutes): the LGM (as from CHELSA, Karger, Nobis, Normand, Graham & Zimmermann, 2021), the Heinrich Stadial1 (17.0-14.7 kyrs BP), the Bølling-Allerød (14.7-12.9 kyrs BP) the Younger Dryas Stadial (12.9-11.7 kyrs BP) and the Holocene: early-Holocene, Greenlandian (11.7-8.326 kyrs BP), mid-Holocene, Northgrippian (8.326-4.2 ka BP) and late-Holocene, Meghalayan (4.2-0.3 kyrs BP) (as from Fordham et al., 2017).

The *O. aveyronensis* occurrences dataset consists of a set of field observations as well as from records downloaded from the gbif (www.gbif.org) and the iNaturalist (www.inaturalist.org) databases before manual curation. The Ecological Niche Modelling analyses were conducted with the R-package *ENMwizard v.0.3.7* (Heming, Dambros & Gutiérrez, 2018), which is a convenient wrapper of several tools. We first spatially filtered occurrences to keep only those that were at least 5 km away from each other using the R-package *spThin* (Aiello-Lammens, Boria, Radosavljevic, Villela & Anderson, 2015). The calibration area for the models was created as a buffer of 0.5° around the minimum convex polygon encompassing all occurrences. From the 19 bioclimatic variables, we selected the less correlated ones (Pearson correlation coefficient < 0.75) using the R-package *caret* (Kuhn, 2019) and kept seven variables: bio2, bio3, bio8, bio9, bio10 and bio17 (see details in Supplementary Appendix S5) for further analyses.

We used the maximum entropy method (implemented in MaxEnt ver. 3.4.1, Phillips, Anderson & Schapire, 2006; Philips & Dudík, 2008; Phillips et al., 2017) to calibrate models and evaluated models’ performance with the package *ENMeval* (Muscarella et al., 2014) as implemented in *ENMwizard*. We evaluated models using a geographic partition scheme of type “block” and optimized two parameters of MaxEnt: the Regularization Multipliers (RM) and the Feature Classes (FCs). RM was varied from 0.5 to 4.5, incremented by 0.5 whereas a suite of 15 FCs (L, for Linear, P, for Product, Q, for Quadratic and H for Hinge) or combination of them were evaluated: L, P, Q, H, LP, LQ, LH, PQ, PH, QH, LPQ, LPH, LQH, PQH, LPQH, resulting in a total of 135 models. Model selection was done by computing the corrected Akaike Information Criterion (“LowAIC”). Model accuracy was also evaluated by calculating omission rates calculated when a 10^th^ percentile threshold is applied (x10ptp). The final model was projected on current and paleoclimatic contexts.

## Results

### Population genomic dataset

We obtained a total of 150 694 350 of read pairs across the 86 individuals (1 382 768 - 7 370 366 of raw reads per ind., mean 3 504 519, SD = 1 282011) of which 290 431 210 barcoded reads (*i.e.* 96.36%) were retained, 10 819 014 (3.6%) were removed because RAD cutsite was not found in the sequence and 138 476 reads (< 1%) were removed because of low quality. We genotyped a total of 859 035 *loci* (composed of 135 255 870 sites) including 368 712 variant sites (SNPs) with *Stacks*. Average read depth ranged from 24.1X to 81.6X (mean 48.7X, SD = 12.4X) based on the combination of parameters we used. After filtering the data with *populations*, we finally kept 4302 *loci* from which we retained 9301 variant sites (SNPs).

### Genetic diversity, population differentiation and divergence

The PCA biplot shows that individuals are arranged by subspecies/country of origin along PC1 (which explains 8.50% of the total genetic variance) and by sampling site along PC2 (which explains 3.88% of the total genetic variance) even though the sampling sites of 1- Guilhaumard and 2-Lapanouse-de-Cernon slightly overlap (Fig. 2a). Based on PC2, intra-population genetic variation seems minimal for 1-Guilhaumard and 2-Lapanouse-de-Cernon and maximal for 3-Saint-Affrique and 5-Larraona. Basic population genetics statistics are indicated in Table 1. All populations displayed weak but significant deviation from panmixia with a mean *G*_IS_ = 0.060 (*p* < 0.01, 95% CI: 0.056-0.064) and population *G*_IS_ values ranging from 0.006 in 3-Saint-Affrique to 0.081 in 4-Valgañón (all *p* < 0.01). Average nucleotide diversity (*π*) varies from 0.269 in 3-Saint-Affrique to 0.290 in 4-Valgañón and 5-Larraona, and allelic richness (*A*_R_) from 16696 (in 3-Saint-Affrique) to 17297 (in 5-Larraona). Based on the filtering procedure we followed, the number of private alleles was relatively low but consistently higher for *O*. *a*. subsp. *vitorica* (*i.e.* from 9 to 23) compared to *O*. *a*. subsp. *aveyronensis* (*i.e.* from 0 to 2).

**Figure 2.**
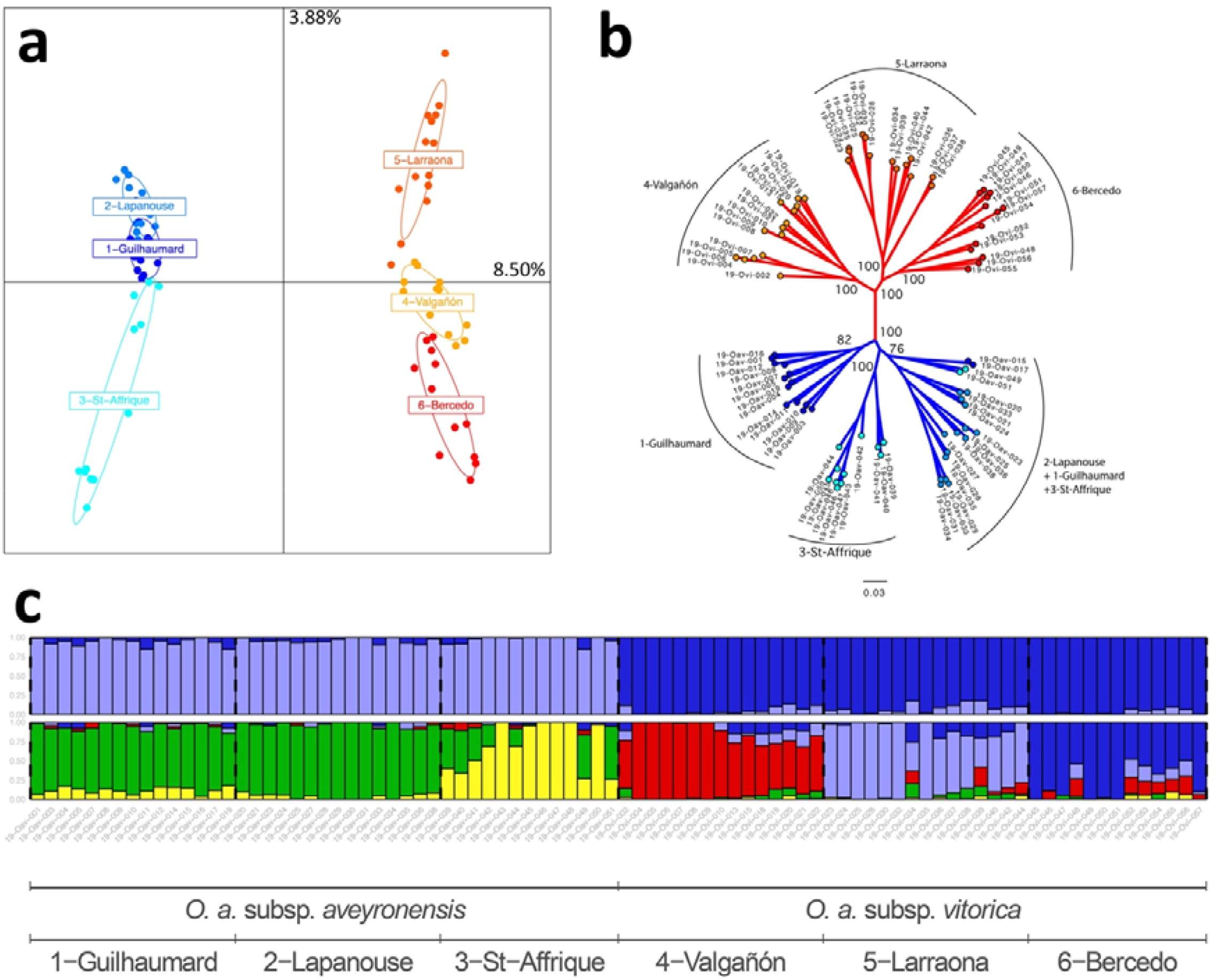
(a) Principal Component Analysis (PCA) displaying the two first axes (PC1 and PC2) representing 8.50% and 3.88% of the total genetic variance. PCA was computed based on 86 individuals genotyped at 9301 SNP and colors depict sampling localities. (b) Maximum Likelihood phylogenetic tree (rooted at midpoint) obtained from the concatenation of the SNPs (node support for main nodes are given as UltraFast Boostrap Approximation based on 1000 replicates). (c) Barplots of ancestry coefficients obtained from sNMF for 86 individuals for *K* = 2 and *K* = 5.

Overall *G*-statistics show that the whole set of populations is genetically structured: *G*_ST_ = 0.091 (*p* < 0.01, 95% CI: 0.089-0.093); *G*^’^_ST_ = 0.134 (*p* < 0.01, 95% CI: 0.105-0.011); *G*’’_ST_ = 0.150 (*p* < 0.01, 95% CI: 0.146-0.153) and Jost’s *D* = 0.047 (*p* < 0.01, 95% CI: 0.046-0.048).

The AMOVA confirms that significant proportions of the total genetic variance are found among populations within subspecies/country (6.8%, *F*_SC_ = 0.072, 95% CI: 0.070-0.074) and between subspecies/country (6.1%, *F*_CT_ = 0.061, *F*_CT_ = 0.058-0.064). Pairwise differentiation between sampling sites varies from *F*_ST_ = 0.049 to 0.082 within subspecies/country and reaches *F*_ST_ = 0.111 to 0.147 between subspecies/country of origin (all *p* < 0.01; see Supplementary Appendix S3 for details).

The clustering analysis carried out with sNMF suggests an optimal number of genetic clusters of *K*=5 based on the cross-entropy criterion (Fig. 2c). We plotted the *Q*-matrices (that contain individual admixture coefficients) corresponding to the best replicate runs for *K*=2 (to investigate whether the program could distinguish the two subspecies) and *K*=5 (Supplementary Appendix S3). For *K*=2, the individuals of *O*. *a*. subsp. *aveyronensis* and *O*. *a*. subsp. *vitorica* were unambiguously assigned to two distinct clusters. For the most likely combination of *K*=5, we observe one cluster per sampling site except for the geographically close localities of 1-Guilhaumard and 2-Lapanouse-de-Cernon that were grouped together in a unique cluster. In the locality of 3-Saint-Affrique 4 out of 13 individuals (19-Oav-039, 19- Oav-040, 19-Oav-049 and 19-Oav-051) display >50% of ancestry proportion that are associated to the cluster associated with 1-Guilhaumard and 2-Lapanouse-de-Cernon.

The phylogenetic tree inferred from the SNP matrix corroborates these results. *O. a*. subsp. *aveyronensis* and *O. a.* subsp. *vitorica* appeared as two well supported reciprocally monophyletic groups (Fig. 2b). Sub-clades corresponding to sampling sites also confirm a phylogenetic sub-clustering consistent with geography in each country for each subspecies (with the exception of a group consisting of a mixture of individuals of all sites in France).

### Inference of past demographic history

Reconstruction of historical dynamics of effective population size using STAIRWAYPLOT2 shows congruent trends consisting of progressive >95% decline in estimated *N*_e_ for all 6 populations, since the LGM, or at least, over the last 25000 years (see Fig. 3). Current effective population sizes were estimated to vary between 302 (2-Lapanouse-de-Cernon) and 642 individuals (3-St-Affrique) with this method. Despite the current geographical discontinuity between *O*. *a*. subsp. *aveyronensis* and *O*. *a*. subsp. *vitorica* populations, we attempted to statistically compare demographic scenarios with and without ongoing migration using DILS. The Posterior Probabilities associated with the models of ongoing migration (*i.e.* IM and SC) relative to models without (*i.e.* SI and AM) was 0.59 based on the pooled dataset and ranged from 0.55 to 0.66 based on the population pairs analyses. These probabilities are low and lead to ambiguous model choices. Therefore, we cannot draw strong conclusions from model comparisons with DILS (see Table 2, Fig. 4 and Appendix S4). Ambiguous inferences can happen when the separation time between the two subspecies (here, *O*. *a*. subsp. *aveyronensis* and *O*. *a*. subsp. *vitorica*) is too recent compared to their current population sizes (Burban, Tenaillon & Glémin, 2024). The median of the parameters estimated based on posterior distributions made under the most probable model (*i.e.* Isolation-with-Migration, IM) indicate very low migration rates (*N.m* = 0.60 for the pooled dataset, and it ranges from 0.56 to 0.92 based on population pairs). With the pooled dataset, we inferred a subdivision of the ancestral population of about 75 000 individuals based on two pools (90 000 to 103 200 based on population pairs) into two daughter populations about 1500 generations ago (650 to 3880 based on population pairs), which convert into 4500 years assuming a generation time of 3 years, 7500 years assuming a generation time of 5 years and 10500 years assuming a generation time of 7 years. These estimates of splitting times were found to be even more recent based on the population pairs (*i.e.* namely, sampling sites) data set. This suggests that most coalescence events are taking place in the ancestral population and not in the respective daughter populations. In other words, the amount of shared polymorphism is still relatively high because of insufficient time for drift to differentially fix/eliminate alleles within the two subranges (*i.e.* Incomplete Lineage Sorting). Similar to what was inferred with STAIWAYPLOT2, our analyses also point to a decrease of effective sizes compared to the ancestral population (*i.e.* current *N*_e_ have values corresponding to about 10% of *N*_anc_). The goodness-of-fit test indicates that the best estimated scenario (IM) reproduces the observed data set with high fidelity (see Supplementary Appendix S4).

**Figure 3.**
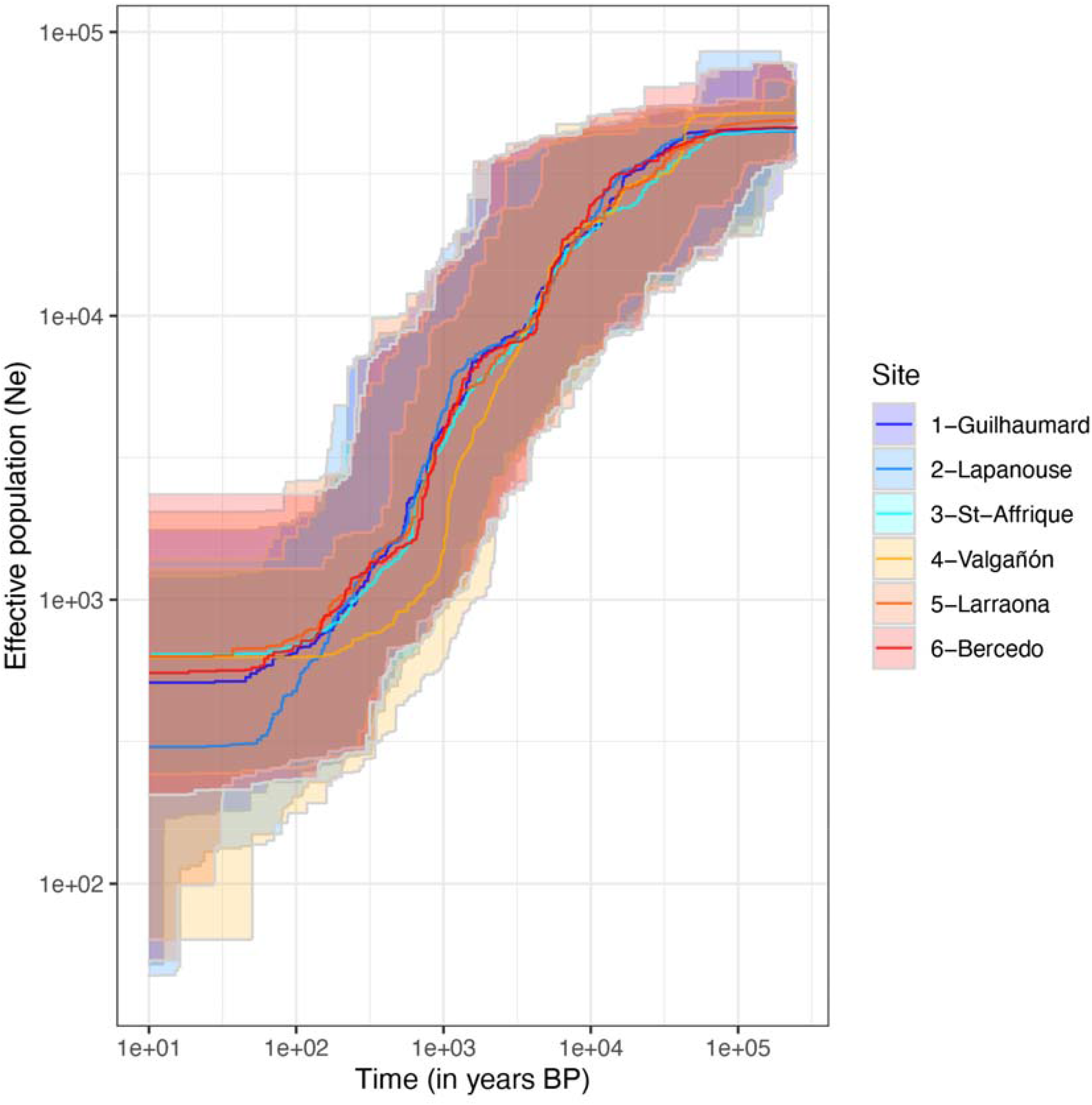
Demographic history of the 6 studied populations of *O. a.* subsp. *aveyronensis* (shades of blue) and *O. a.* subsp. *vitorica* (shades of red) depicting the mean effective population size *N*_e_ (and associated 95 confidence intervals) over time as inferred from STAIRWAYPLOTv2, assuming a mutation rate of 7 x10^-9^ and a generation time of 5 years.

**Figure 4.**
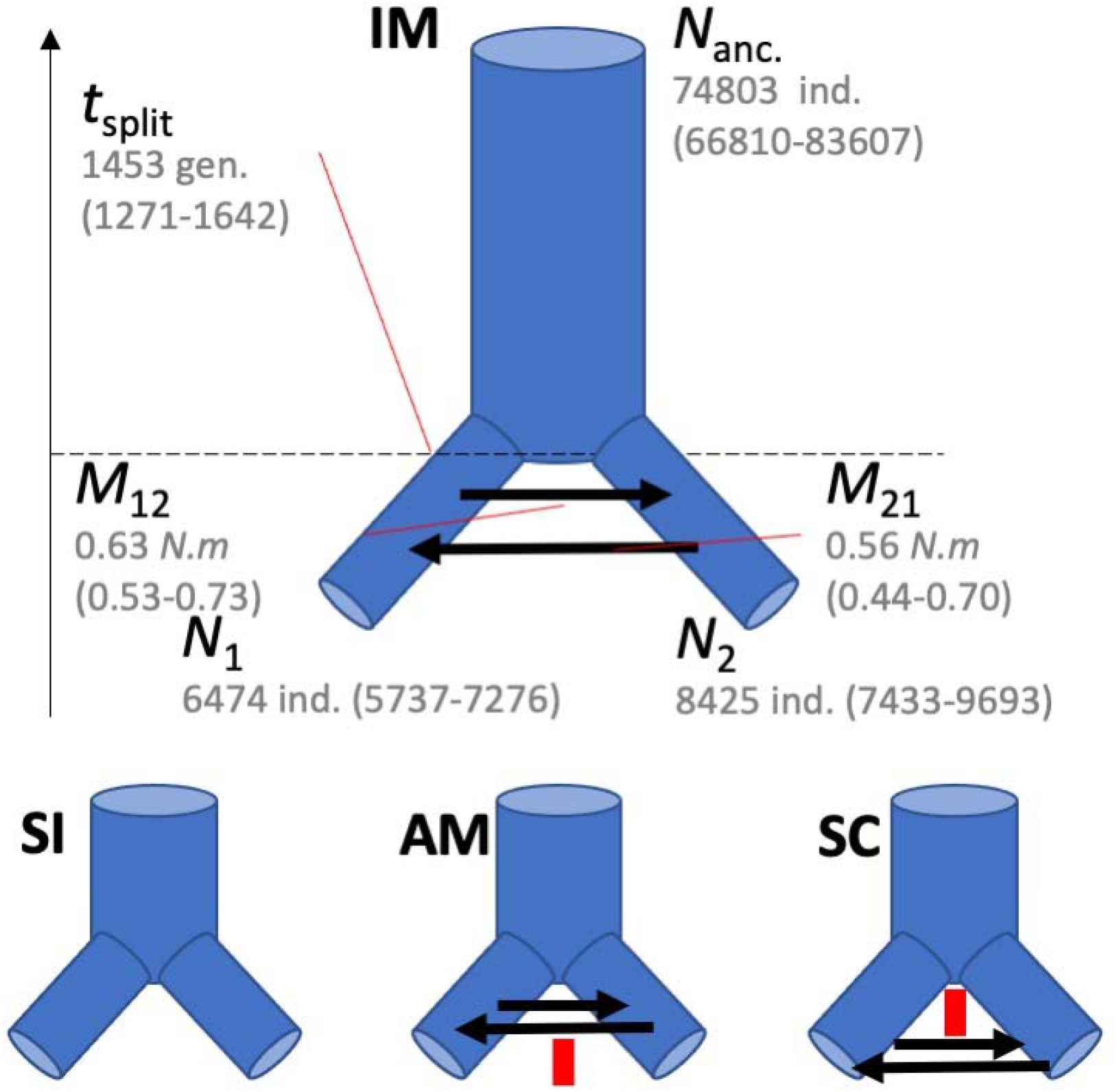
Schematic overview of the best demographic scenario (Isolation with Migration, IM) for pooled dataset (*i.e. O. aveyronensis* subsp. *aveyronensis*/ *O. a.* subsp. *vitorica*). The median values (and 2.5-97.5% boundaries of Highest Posterior Distribution probability) are shown for the parameters inferred with DILS. *T_split_*: time of split (given in number of generations), *N*_a_: ancestral effective population size, *N*_1_: (current) effective population size for the *O. a.* subsp. *aveyronensis* population (given in number of individuals), *N*_2_: (current) effective population size for the *O. a.* subsp. *vitorica*, *M*_12_: migration rate from *O. a.* subsp. *aveyronensis* to *O. a.* subsp. *vitorica* and *M*_21_: migration rate from *O. a.* subsp. *vitorica* to *O. a.* subsp. *aveyronensis*. Alternative demographic scenarios tested are also illustrated: Strict Isolation (SI), Ancestral Migration (AM, involving a cessation of gene flow after an initial stage of migration after the split) and Secondary Contact (SC, involving a secondary contact after an initial stage of isolation after the split).

**Table 2.** Summary of the main demographic parameters inferred with DILS i) for pooled dataset (*Ophrys aveyronensis* subsp. *aveyronensis* and *<u>Ophrys aveyronensis</u>* subsp. *vitorica*) and ii) for each *Ophrys aveyronensis* subsp. *aveyronensis* and *Ophrys aveyronensis* subsp. *vitorica* population pairs. *N*_e_ 1, *N*_e_ 2, *N*_e_ anc. correspond to effective population sizes estimated for population 1, population 2 and the ancestral population, *t*_split_ to the time of split between population 1 and population 2 in number of generations, respectively and M_12_/M_21_ the migration rates (as *N*_e_.m) from population 1 to population 2 and from population 2 to population 1. Parameters have been estimated based on a model using neural network.

| Population 1 | Population 2 | $N_e$ 1 | $N_e$ 2 | $N_e$ anc. | $t_{split}$ | $M_{12}/M_{21}$ |
| --- | --- | --- | --- | --- | --- | --- |
| O. a. *aveyronensis* pool (France) | O. a. *vitorica* pool (Spain) | 6474 (5737-7276) | 8425 (7433-9693) | 74803 (66810-83607) | 1453 (1271-1642) | 0.63/0.56 (0.53-0.73/0.44-0.70) |
| 1-Guilhaumard | 4-Valgañón | 4934 (4290-5662) | 4105 (3622-4589) | 98474 (92517-105003) | 1226 (1079-1383) | 0.90/0.74 (0.73-1.07/0.60-0.88) |
| 1-Guilhaumard | 5-Larraona | 2403 (2089-2751) | 2968 (2569-3429) | 103206 (97489-109542) | 651 (573-724) | 0.89/0.64 (0.72-1.06/0.52-0.76) |
| 1-Guilhaumard | 6-Bercedo | 9824 (8553-11446) | 10477 (7866-13078) | 97808 (93302-103547) | 3029 (2672-3410) | 0.73/0.44 (0.35-0.53/0.60-0.86) |
| 2-Lapanouse | 4-Valgañón | 2729 (2506-3035) | 2640 (2311-2953) | 99913 (94450-105541) | 803 (728-872) | 0.46/0.67 (0.38-0.55/0.54-0.79) |
| 2-Lapanouse | 5-Larraona | 4061 (3556-4554) | 6049 (5291-6972) | 93345 (89784-97211) | 1416 (1255-1564) | 0.42/1.01 (0.34-0.50/0.71-1.28) |
| 2-Lapanouse | 6-Bercedo | 5812 (4988-6822) | 7719 (6431-9262) | 97785 (90460-105961) | 2069 (1731-2410) | 0.84/0.74 (0.67-1.00/0.58-0.93) |
| 3-St-Affrique | 4-Valgañón | 9995 (8557-11455) | 13510 (11837-15468) | 94633 (89865-100052) | 3879 (3236-4549) | 0.89/0.81 (0.71-1.05/0.62-0.99) |
| 3-St-Affrique | 5-Larraona | 7270 (6386-8230) | 10997 (9603-12667) | 90012 (86305-93995) | 2891 (2494-3300) | 0.82/0.79 (0.66-0.97/0.56-0.98) |
| 3-St-Affrique | 6-Bercedo | 9562 (8393-10964) | 10997 (9632-12459) | 99195 (92748-104706) | 3682 (3184-4204) | 0.95 /0.89 (0.77-1.14-0.69-1.08) |

### Current and past Species Distribution Modelling

ENMeval analyses identified RM = 1 and a mixture of Product (P), Quadratic (Q) and Hinge (H) feature classes as the best-performing parameters for calibrating the final ENM. These parameters yielded a single “best” candidate model with the lowest AICc score (925.02) and an AICc weight of ∼0.23. This model had mean omission rates of 0.165 (for the 10^th^ percentile) and a mean test AUC of 0.834 (see variable contributions following this model in Table 3 and Supplementary Appendix S5, for additional details). The spatial projection of the current bioclimatic niches inferred for *O. aveyronensis* corresponds to its actual geographic distribution but show that some additional areas may be climatically suitable for this orchid (Fig. 5). Spatial projections considering paleoclimatic conditions show that the extent of suitable habitat for *O. aveyronensis* may have reached its apex during the Younger Dryas cold period (12.9-11.7 kyrs BP) before starting to decrease across the Holocene. The LGM conditions (21 kyrs BP) were not suitable for the establishment of viable populations in the studied area but important suitable habitats may have existed during the Heinrich Stadial 1 event (17-14.7 kyrs BP) and the Bøllering-Allerød warming (14.7 – 12.9 kyrs BP). Altogether, these results are consistent with the possible existence of a unique ancestral area, centered on South-Western France that could have experienced a split and a combination of reduction and shift towards cooler mid-elevation areas, in the North-East (for *O. a.* subsp. *aveyronensis*) and in the South-West (for *O. a.* subsp. *vitorica*) during the Holocene. The results available from WorldClim data do not contradict these conclusions (Appendix S5). However, the latter results show significant differences across GCMs and even for a given GCM but downscaled based on different interpolation algorithms: namely, WorldClim instead of CHELSA.

**Figure 5.**
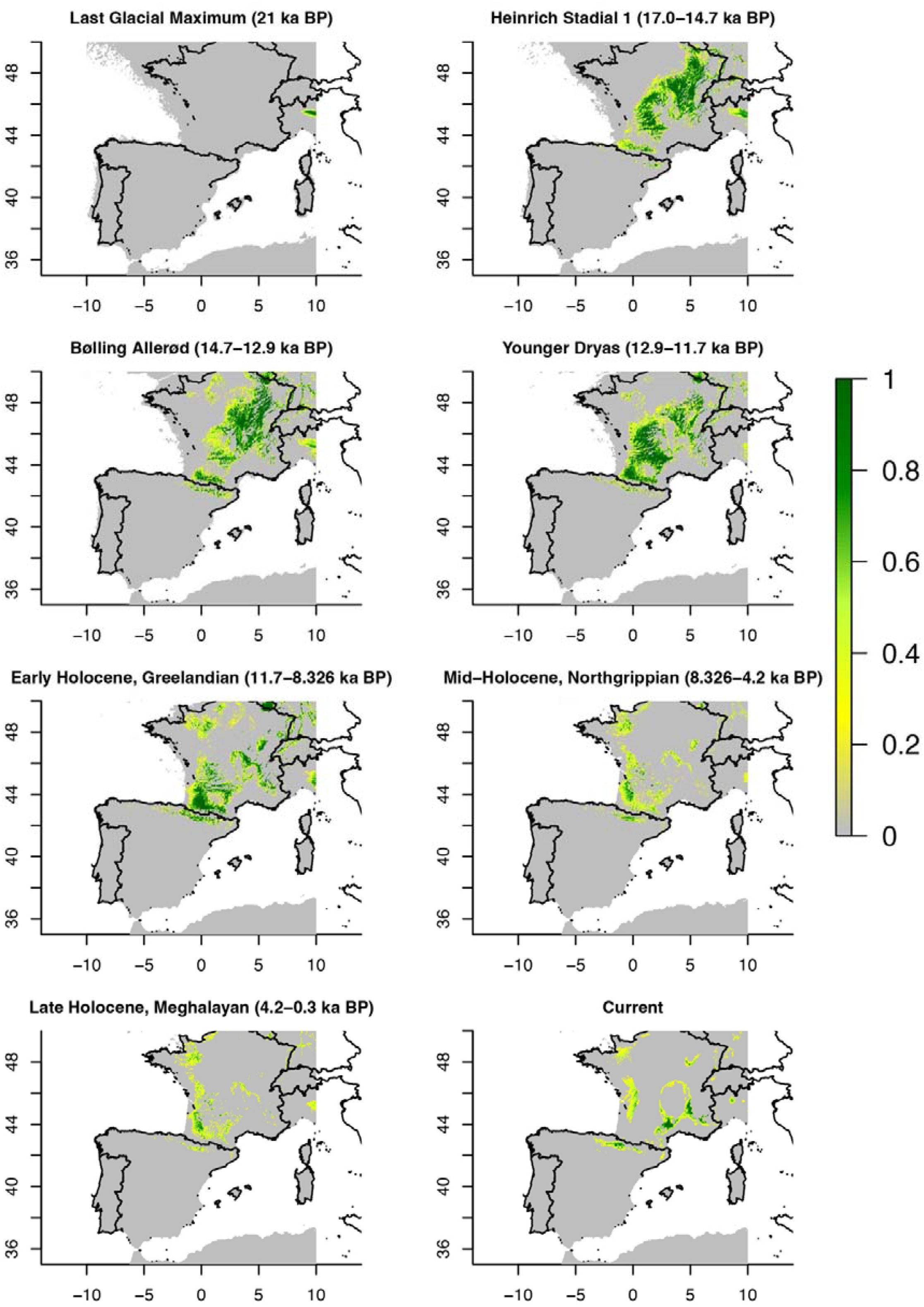
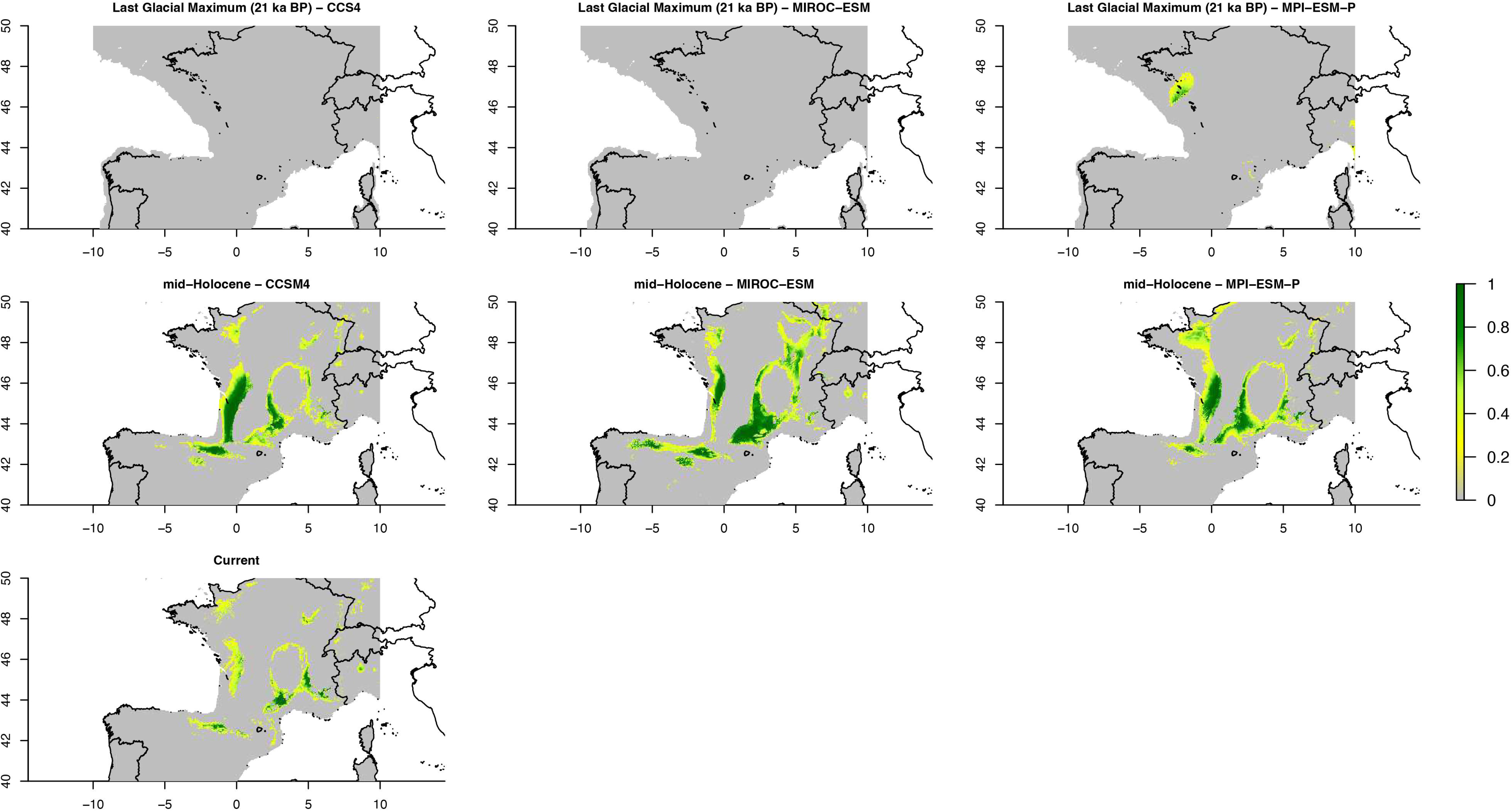
Spatial projections of the bioclimatic niches inferred for the *Ophrys aveyronensis* depicting habitat suitability at different time periods from the Last Glacial Maximum (LGM, about 21 kyrs ago) and with projections for the Heinrich Stadial1 (17.0-14.7 kyrs BP), the Bølling-Allerød (14.7-12.9 kyrs BP) the Younger Dryas Stadial (12.9-11.7 kyrs BP) and the Holocene: early-Holocene, Greenlandian (11.7-8.326 kyrs BP), mid-Holocene, Northgrippian (8.326-4.2 kyrs BP) and late-Holocene, Meghalayan (4.2-0.3 kyrs BP) and currently.

**Table 3.** List of bioclimatic variables available from WorldClim. From these 19 variables (bio1 to bio19) the ones used for ENM analyses (Person correlation coefficient < 0.75) are indicated in bold with their relative contribution in the selected ENM model.

| Variable name | Denomination | Relative contribution (in %) |
| --- | --- | --- |
| bio1 | Annual Mean Temperature |  |
| **bio2** | **Mean Diurnal Range (Mean of monthly (max temp - min temp))** | **8.14** |
| **bio3** | **Isothermality (bio2/bio7) (×100)** | **5.32** |
| bio4 | Temperature Seasonality (standard deviation ×100) |  |
| bio5 | Max Temperature of Warmest Month |  |
| bio6 | Min Temperature of Coldest Month |  |
| bio7 | Temperature Annual Range (bio5-bio6) |  |
| **bio8** | **Mean Temperature of Wettest Quarter** | **3.21** |
| **bio9** | **Mean Temperature of Driest Quarter** | **3.01** |
| **bio10** | **Mean Temperature of Warmest Quarter** | **11.31** |
| bio11 | Mean Temperature of Coldest Quarter |  |
| bio12 | Annual Precipitation |  |
| bio13 | Precipitation of Wettest Month |  |
| bio14 | Precipitation of Driest Month |  |
| **bio15** | **Precipitation Seasonality (Coefficient of Variation)** | **41.65** |
| bio16 | Precipitation of Wettest Quarter |  |
| **bio17** | **Precipitation of Driest Quarter** | **23.34** |
| bio18 | Precipitation of Warmest Quarter |  |
| bio19 | Precipitation of Coldest Quarter |  |

## Discussion

Some *Ophrys* orchid lineages (e.g. *O. sphegodes sensu lato*) boast some of the highest diversification rates reported worldwide for orchids and angiosperms in general (Breitkopf et al., 2015; see also Thompson, Davis, Doff, Willis & Priest, 2023) and are being increasingly mentioned as biological models of interest for studying ecological speciation and adaptive radiations (e.g. Nosil et al., 2017; Baguette et al., 2020; Nürk et al., 2020). Ecological and sometimes sympatric speciation in *Ophrys* is confidently explained by the selective pressures exerted by pollinators and the high degree of specificity required to ensure the success of the sexual swindling strategy (see Baguette et al., 2020). Nevertheless, allopatric-vicariant divergence undoubtedly also plays an important role in the diversification of *Ophrys*. In this study, we used population genomics and ENM to characterise one of the most recent cases of allopatric-vicariant divergence for a continental plant. *Ophrys aveyronensis* therefore provides a promising opportunity to investigate the relative contribution of biogeographic and ecological factors in shaping early stages of evolutionary divergence in rapidly diversifying groups of organisms.

### Patterns of genetic diversity and differentiation based on population genomics

Although populations of *Ophrys aveyronensis* from southern France (subsp. *aveyronensis*) and northern Spain (subsp. *vitorica*) differ only in subtle ways on the basis of phenotypic data (Gibert et al., 2022; 2024), our results allowed to unambiguously tease them apart based on population genomics. Both the PCA, the *F*-statistics (and/or their analogues), the clustering analysis and the phylogenetic tree confirm that the two subspecies form distinct genetic entities, which themselves show a lower genetic substructure between sampling sites within subranges. The reciprocal monophyly of the two disjunct subspecies (with no outlier individuals deviating from the global pattern) *a priori* rules out a recent dispersal scenario. We found that all the populations sampled had similar levels of allelic richness, heterozygosity, departure from panmixia and number of private alleles. There is therefore no evidence that any of these six populations represent a subset of the genetic diversity of a putative source population elsewhere. Finally, the inference of the temporal dynamics of effective population size suggests that each of the sampled populations can be considered as a kind of replicate of each other. All the populations displayed consistent decline in effective population size over the last tens of thousands of years, suggesting that they shared a similar demographic history. In summary, these results are fully consistent with a scenario of range splitting, and subsequent contraction of two subranges with restricted inter-regional gene flow.

Delforge (2016) hypothesised that the Iberian populations could be the result of a hybrid swarm (without mentioning the putative parental species), undergoing spatial and demographic expansion. According to this scenario, the two morphologically similar population groups may have evolved convergent phenotypes as a result of their adaptation to the same insect pollinator species on both sides of the Pyrenees. Such evidence of phenotypic convergence between *Ophrys* species originating from different phylogenetic lineages has been already documented (Stökl et al., 2005; Gögler et al., 2009; Sramkó, Gulyas & Molnár, 2010). However, without ruling out the possibility that both subspecies have experienced gene flow with other sympatric *Ophrys* taxa, the striking similarity we found in patterns of genetic diversity, as well as in the temporal trends we inferred for the effective sizes of all populations sampled in this study (regardless their geographic origin) do not support this hypothesis, but rather suggest that *O. a.* subsp. *aveyronensis* and *O. a.* subsp. *vitorica* originated from a single and direct common ancestor.

The demographic scenarios we tested remain relatively simple and the Bayesian posterior probabilities we obtained with DILS show that the approach implemented in this program, while favouring ongoing migration, could not strongly support such a model over isolation ones. A theoretical explanation for this otherwise puzzling result is that the time of split is too recent to observe sufficient allele sorting (and signal) from a diverse ancestral population with relatively high effective population size to be able to detect a cessation of gene flow between the two species (see Smith & Hahn, 2023, Burban et al. 2024). In addition, events that are known to deceive demographic inferences such as ‘ghost introgression’ from unsampled lineages (Tricou, Tannier & De Vienne, 2022) may have a non-negligible probability to occur in *Ophrys*. Investigating whether or not *Ophrys aveyronensis* has a hybrid origin, and at least whether and to what extent the two taxa have been introgressed by other *Ophrys* taxa will require the genomic analyses of additional species in a future study. Nevertheless, we are confident that the time of split we inferred here is among the rare cases documenting a so recent speciation event in a plant continental system (see de Queiroz et al., 2022), given that ghost introgression may have led to an overestimation of the time of split (Pang & Zang 2023).

### On the causes of the disjunct geographic distribution of O. aveyronensis

The results of the Ecological Niche Modelling show that the conditions of the LGM were not suitable for the establishment of viable populations of *O. aveyronensis* . This suggests that either this species diversified from an ancestral *Ophrys* lineage in the study area after this period, or that its populations occupied a relatively remote glacial refugium during this period (the ENM results suggest that the Italian peninsula could have provided such a refugium, see Fig. 5). During the last 21000 years, the extent of suitable habitat for *O. aveyronensis* was rather large until the Holocene (about 12000 years ago) (Fig. 5), when its area began to decrease dramatically while maintaining an area of relatively high suitability centred on southwestern France. The split and, to a lesser extent, the northward and southward shifts may have occurred as late as during the Late Holocene (from 4000 years ago) (Fig. 5). Interestingly, this timing is the same order of magnitude as the time of the split we inferred from the population genomic data, and is consistent with a scenario of allopatric divergence (without being able to completely exclude transient gene flow through climate-induced oscillations throughout the Holocene) from an ancestral population with relatively large effective size.

The Pyrenees mountains have the peculiarity of acting as a biogeographic boundary for many terrestrial organisms, while being at the crossroads between temperate and Mediterranean biomes (see Pironon, Gómez, Font & García, 2022 and references therein). Although orchid propagules have the capacity to disperse efficiently over long distances, several taxa occur on either the southern or the northern slopes of the Pyrenees, but have ranges that do not extend latitudinally on either side, or only weakly so (*i.e.* the so-called northern or southern peripheral species in Pironon et al., 2022). The Pyrenees may therefore alter the dispersal of at least some orchid species and populations (e.g. *Dactylorhiza insularis*, *Gymnadenia gabasiana*, *Neotinea conica*, *Ophrys castellana*, *O. riojana* and *O. subinsectifera*, *Orchis langei* and *O. simia*) due to various biotic or abiotic factors. This mostly granitic and relatively high mountain channel forms a vast area of unsuitable habitat for *Ophrys* orchids, which generally require calcareous soils and lowland or midland environments. Although data on their geographical distribution remain incomplete, we may also speculate that the absence of *Ophrys*-specific pollinator species and/or of fungi on which the orchid depends for their germination and their development may also have limited the possibility of establishment of intermediate orchid populations between the two current subranges. For *O. aveyronensis*, our results are consistent with a general reduction in gene flow between both sides of the Pyrenees through the formation and the contraction of the two subranges during the Holocene until the present cessation of gene exchange, which is too recent to be supported on the basis of the model-based demographic inferences but is corroborated by our descriptive genetic analyses.

### Evidence for ecological speciation in addition to vicariance?

Selection pressures exerted by insect pollinators are likely to be the main driver of ecological and presumably sympatric speciation in *Ophrys* orchids (see Baguette et al., 2020). Thus, populations of a given *Ophrys* species that are pollinated by distinct insect pollinator species can be considered to be undergoing a speciation process. So far, field observations support that *O. aveyronensis* from the Grands Causses and their Iberian counterparts are pollinated by the same pollinator species: the solitary bee *Andrena hattorfiana* (see Benito Ayuso, 2019). However, in a recent study on the same set of individuals, we showed that *O. aveyronensis* from the Grands Causses and from the Iberian Peninsula show subtle differences in a number of morphological and colour traits that may be involved in pollination (Gibert et al., 2022; 2024). These subtle differences may arise as a result of local adaptation to a “secondary” pollinator species (see Joffard et al., 2019; Schatz, Genoud, Claessens & Kleynen, 2020; Baguette et al., 2021). Ongoing work is now addressing whether this variation is indeed actually adaptive and involved in evolutionary divergence i) by assessing which phenotypic traits may be under divergent selection among the *O. aveyronensis* populations across their range and which ones may be evolving ‘non-ecologically’ and ii) by investigating putative genomic regions associated with phenotypic trait values, genetic structure or ecological factors, something that remains largely underexplored in *Ophrys* so far (but see Gibert et al., 2024).

Cross-pollination experiments between individuals of the two subranges could also help to highlight the insurgence of post-zygotic reproductive isolation in *Ophrys aveyronensis*. However, such work may be hampered by the difficulty of growing *Ophrys* under controlled conditions and by the protection status of *Ophrys aveyronensis* in France. Current knowledge suggests that such reproductive barriers may be weak or absent at the scale *Ophrys* genus (Soliva and Widmer, 2003; Scopece, Mesacchio, Widmer & Cozzolino, 2007, Gervasi et al., 2017). To our best knowledge, only two notable exceptions to this pattern involve the phylogenetically distant *O. incubacea* and *O. iricolor* (Cortis et al., 2009) or species with different ploidy levels, such as the diploid *O. exaltata* and the tetraploid *O. lupercalis* (Vereecken, Cozzolino & Schiestl, 2010).

### Taxonomic and conservation implications

Given that populations of *O. aveyronensis* are pollinated by the same insect pollinator species throughout their range and have similar environmental niches, we currently have no evidence to even consider the existence of distinct ecotypes in *Ophrys aveyronensis* . However, the disjunct geographic distribution in two subranges, separated by ∼600 km and the Pyrenees Mountains suggests that populations from the Grands Causses and from the Iberian Peninsula may be considered as distinct evolutionary (and conservation) units. Our population genomic analyses suggest that both sets of populations are genetically differentiated and are unlikely to have experienced ongoing gene flow. Accordingly, we propose that they should be considered as two subspecies: *Ophrys aveyronensis* subsp. *aveyronensis* and *Ophrys aveyronensis* subsp. *vitorica*, following Paulus (2017).

Recent studies have elaborated on some empirical evidence that the delineation of the so-called ‘micro-species’ in *Ophrys* is practically cumbersome (Bateman et al., 2021; Bateman and Rudall, 2023), especially in the section *Sphegodes* to which *O. aveyronensis* belongs. Some genomic data, such as whole plastomes, which are useful to delineate the main *Ophrys lineages*, show a sequence similarity degree of more than 99.5% between ‘microspecies’ and therefore do not show enough variation to allow to distinguish them (Bertrand, Gibert, Llauro & Panaud, 2019; Bertrand Gibert, Llauro & Panaud, 2021; Bateman et al., 2021). Here, however, our results demonstrate that an approach consisting of sampling populations (*i.e.* a sufficient number of individuals at a given location) and appropriate genetic markers (e.g. SNPs genotyped on populations based on RAD-seq protocols) provides signals that are consistent with taxonomy. We therefore argue that such approaches of population genomics within an integrative taxonomy framework (relying on flower morphology, pollinator identity, ecological features…) is by far the most fruitful for better understanding the process of evolutionary divergence and, at the same time, for improving our knowledge of the taxonomy, systematics, ecology and evolution of *Ophrys* orchids.

### Ophrys aveyronensis as a promising biological model to study speciation

Vicariance is a very common route toward speciation (Hewitt et al., 2000), but most of empirical cases document it retrospectively, between nascent species that exhibit at the same time obvious geographic separation and substantial levels of both phenotypic and genetic differentiation (Dool, Picker & Eberhard, 2021). In this context, studies reporting cases of incipient allopatric divergence are rare in plants, especially in continental systems (see de Queiroz et al., 2022). The allopatric divergence in the *Ophrys aveyronensis* we report here is indeed extremely recent compared to other species with disjunct geographic distributions in terrestrial systems (e.g. González-Serna, Cordero & Orego, 2018; Tomasello, Karbstein, Hodač, Paetzold & Hörandl, 2020; Wang et al., 2022; Cao et al., 2022; Balmori-de la Puente et al., 2022). It is likely to have arisen as a result of climate warming, not as a consequence of Quaternary climate fluctuations but as late as following the Last Glacial Maximum, which disrupted an ancestral geographic distribution and subsequently constrained the spatial dynamics of its subranges and does not *a priori* imply a human-induced biological introduction. This system may therefore serve as a natural laboratory to study parallel adaptation at early stages of reproductive isolation insurgence, following climate-induced range fragmentation.

### Data availability statement

Sequencing data have been submitted to the European Nucleotide Archive (ENA; https://www.ebi.ac.uk/ena/browser/home) under Study with primary accession n°PRJEB61037 (and secondary accession number ERP146119) and samples accession n°ERS14864169 (SAMEA112857509) to n°ERS14864254 (SAMEA112857594).

## Author contributions

JB, BS, AG and MB designed the study. AG, JB, BS and RB gathered the data. AG and JB performed the analyses with contribution from CF and CR and wrote the first version of the manuscript. All authors contributed substantially to the revisions.

## Acknowledgements

This work was primarily supported by an inter-LabEX TULIP-CEMEB initiative (n°197310) to J. Bertrand and B. Schatz. This research was also funded by an Agence Nationale pour la Recherche Jeune Chercheur Jeune Chercheuse (ANR JCJC) grant to J. Bertrand, grant number ANR-21-CE02-0022-01, and is set within the framework of the “Laboratoires d’Excellences (LABEX)” TULIP [ANR-10- LABX-41]. Lastly, this research was also funded by the Observatoire de REcherche Méditerranéen de l’Environnement (SO Ocove & SO PolliMed, OSU OREME) to B. Schatz. We thank J.-L. Roux and D. Vizcaïno for the information provided about sampling sites in Spain. We thank C. Moliné for his contribution to fieldwork, M. El Baidouri and P. Salvado for their support with bioinformatic analyses. P. Comes, M. James and two anonymous reviewers also provided constructive feedback on earlier versions of the manuscript.

## Biosketch

A.G. and the team are interested in plant ecology and evolution and in particular on the eco- evolutionary significance of phenotypic variation at early stages of the speciation process. Our aim is to better integrate phenotypic, genotypic and ultimately individual fitness data to understand the onset of reproductive isolation by investigating both its molecular bases and its consequences on phenotypes and fitness. We investigate natural populations of bee orchids (*Ophrys*) and its remarkable insect-mimicking flowers as a model.

