## Supplementary Materials for "Holocene climate change promoted allopatric divergence and disjunct geographic distribution in a bee orchid species"

**Supplementary Table S1** Individual ID, country, population, and collector names.

| Individual ID | Country | Population | Collectors |
| --- | --- | --- | --- |
| 19-Oav-001 | France | Guilhaumard | JB, AG, BS |
| 19-Oav-002 | France | Guilhaumard | JB, AG, BS |
| 19-Oav-003 | France | Guilhaumard | JB, AG, BS |
| 19-Oav-004 | France | Guilhaumard | JB, AG, BS |
| 19-Oav-005 | France | Guilhaumard | JB, AG, BS |
| 19-Oav-006 | France | Guilhaumard | JB, AG, BS |
| 19-Oav-007 | France | Guilhaumard | JB, AG, BS |
| 19-Oav-008 | France | Guilhaumard | JB, AG, BS |
| 19-Oav-009 | France | Guilhaumard | JB, AG, BS |
| 19-Oav-010 | France | Guilhaumard | JB, AG, BS |
| 19-Oav-011 | France | Guilhaumard | JB, AG, BS |
| 19-Oav-012 | France | Guilhaumard | JB, AG, BS |
| 19-Oav-013 | France | Guilhaumard | JB, AG, BS |
| 19-Oav-014 | France | Guilhaumard | JB, AG, BS |
| 19-Oav-015 | France | Guilhaumard | JB, AG, BS |
| 19-Oav-016 | France | Guilhaumard | JB, AG, BS |
| 19-Oav-017 | France | Guilhaumard | JB, AG, BS |
| 19-Oav-018 | France | Guilhaumard | JB, AG, BS |
| 19-Oav-019 | France | Guilhaumard | JB, AG, BS |
| 19-Oav-020 | France | Lapanouse | JB, AG, BS |
| 19-Oav-021 | France | Lapanouse | JB, AG, BS |
| 19-Oav-022 | France | Lapanouse | JB, AG, BS |
| 19-Oav-023 | France | Lapanouse | JB, AG, BS |
| 19-Oav-024 | France | Lapanouse | JB, AG, BS |
| 19-Oav-025 | France | Lapanouse | JB, AG, BS |
| 19-Oav-026 | France | Lapanouse | JB, AG, BS |
| 19-Oav-027 | France | Lapanouse | JB, AG, BS |
| 19-Oav-028 | France | Lapanouse | JB, AG, BS |
| 19-Oav-029 | France | Lapanouse | JB, AG, BS |
| 19-Oav-030 | France | Lapanouse | JB, AG, BS |
| 19-Oav-031 | France | Lapanouse | JB, AG, BS |
| 19-Oav-032 | France | Lapanouse | JB, AG, BS |
| 19-Oav-033 | France | Lapanouse | JB, AG, BS |
| 19-Oav-034 | France | Lapanouse | JB, AG, BS |
| 19-Oav-035 | France | Lapanouse | JB, AG, BS |
| 19-Oav-036 | France | Lapanouse | JB, AG, BS |
| 19-Oav-037 | France | Lapanouse | JB, AG, BS |
| 19-Oav-038 | France | Lapanouse | JB, AG, BS |
| 19-Oav-039 | France | St-Affrique | JB, AG, BS |
| 19-Oav-040 | France | St-Affrique | JB, AG, BS |
| 19-Oav-041 | France | St-Affrique | JB, AG, BS |
| 19-Oav-042 | France | St-Affrique | JB, AG, BS |
| 19-Oav-043 | France | St-Affrique | JB, AG, BS |
| 19-Oav-044 | France | St-Affrique | JB, AG, BS |
| 19-Oav-045 | France | St-Affrique | JB, AG, BS |
| 19-Oav-046 | France | St-Affrique | JB, AG, BS |
| 19-Oav-047 | France | St-Affrique | JB, AG, BS |
| 19-Oav-048 | France | St-Affrique | JB, AG, BS |
| 19-Oav-049 | France | St-Affrique | JB, AG, BS |
| 19-Oav-050 | France | St-Affrique | JB, AG, BS |
| 19-Oav-051 | France | St-Affrique | JB, AG, BS |
| 19-Oav-052 | France | St-Affrique | JB, AG, BS |
| 19-Ovi-001 | Spain | Valgañón | JB, AG, BS, RB, CM |
| 19-Ovi-002 | Spain | Valgañón | JB, AG, BS, RB, CM |
| 19-Ovi-003 | Spain | Valgañón | JB, AG, BS, RB, CM |
| 19-Ovi-004 | Spain | Valgañón | JB, AG, BS, RB, CM |
| 19-Ovi-005 | Spain | Valgañón | JB, AG, BS, RB, CM |
| 19-Ovi-006 | Spain | Valgañón | JB, AG, BS, RB, CM |
| 19-Ovi-007 | Spain | Valgañón | JB, AG, BS, RB, CM |
| 19-Ovi-008 | Spain | Valgañón | JB, AG, BS, RB, CM |
| 19-Ovi-009 | Spain | Valgañón | JB, AG, BS, RB, CM |
| 19-Ovi-010 | Spain | Valgañón | JB, AG, BS, RB, CM |
| 19-Ovi-011 | Spain | Valgañón | JB, AG, BS, RB, CM |
| 19-Ovi-012 | Spain | Valgañón | JB, AG, BS, RB, CM |
| 19-Ovi-013 | Spain | Valgañón | JB, AG, BS, RB, CM |
| 19-Ovi-014 | Spain | Valgañón | JB, AG, BS, RB, CM |
| 19-Ovi-015 | Spain | Valgañón | JB, AG, BS, RB, CM |
| 19-Ovi-016 | Spain | Valgañón | JB, AG, BS, RB, CM |
| 19-Ovi-017 | Spain | Valgañón | JB, AG, BS, RB, CM |
| 19-Ovi-018 | Spain | Valgañón | JB, AG, BS, RB, CM |
| 19-Ovi-019 | Spain | Valgañón | JB, AG, BS, RB, CM |
| 19-Ovi-020 | Spain | Valgañón | JB, AG, BS, RB, CM |
| 19-Ovi-021 | Spain | Valgañón | JB, AG, BS, RB, CM |
| 19-Ovi-022 | Spain | Valgañón | JB, AG, BS, RB, CM |
| 19-Ovi-023 | Spain | Larraona | JB, AG, BS, RB, CM |
| 19-Ovi-024 | Spain | Larraona | JB, AG, BS, RB, CM |
| 19-Ovi-025 | Spain | Larraona | JB, AG, BS, RB, CM |
| 19-Ovi-026 | Spain | Larraona | JB, AG, BS, RB, CM |
| 19-Ovi-027 | Spain | Larraona | JB, AG, BS, RB, CM |
| 19-Ovi-028 | Spain | Larraona | JB, AG, BS, RB, CM |
| 19-Ovi-029 | Spain | Larraona | JB, AG, BS, RB, CM |
| 19-Ovi-030 | Spain | Larraona | JB, AG, BS, RB, CM |
| 19-Ovi-031 | Spain | Larraona | JB, AG, BS, RB, CM |
| 19-Ovi-032 | Spain | Larraona | JB, AG, BS, RB, CM |
| 19-Ovi-033 | Spain | Larraona | JB, AG, BS, RB, CM |
| 19-Ovi-034 | Spain | Larraona | JB, AG, BS, RB, CM |
| 19-Ovi-035 | Spain | Larraona | JB, AG, BS, RB, CM |
| 19-Ovi-036 | Spain | Larraona | JB, AG, BS, RB, CM |
| 19-Ovi-037 | Spain | Larraona | JB, AG, BS, RB, CM |
| 19-Ovi-038 | Spain | Larraona | JB, AG, BS, RB, CM |
| 19-Ovi-039 | Spain | Larraona | JB, AG, BS, RB, CM |
| 19-Ovi-040 | Spain | Larraona | JB, AG, BS, RB, CM |
| 19-Ovi-041 | Spain | Larraona | JB, AG, BS, RB, CM |
| 19-Ovi-042 | Spain | Larraona | JB, AG, BS, RB, CM |
| 19-Ovi-043 | Spain | Larraona | JB, AG, BS, RB, CM |
| 19-Ovi-044 | Spain | Larraona | JB, AG, BS, RB, CM |
| 19-Ovi-045 | Spain | Bercedo | JB, AG |
| 19-Ovi-046 | Spain | Bercedo | JB, AG |
| 19-Ovi-047 | Spain | Bercedo | JB, AG |
| 19-Ovi-048 | Spain | Bercedo | JB, AG |
| 19-Ovi-049 | Spain | Bercedo | JB, AG |
| 19-Ovi-050 | Spain | Bercedo | JB, AG |
| 19-Ovi-051 | Spain | Bercedo | JB, AG |
| 19-Ovi-052 | Spain | Bercedo | JB, AG |
| 19-Ovi-053 | Spain | Bercedo | JB, AG |
| 19-Ovi-054 | Spain | Bercedo | JB, AG |
| 19-Ovi-055 | Spain | Bercedo | JB, AG |
| 19-Ovi-056 | Spain | Bercedo | JB, AG |
| 19-Ovi-057 | Spain | Bercedo | JB, AG |
| Collectors: Joris BERTRAND (JB), Anaïs GIBERT (AG), Bertrand SCHATZ (BS), Roselyne BUSCAIL (RB) and Christian MOLINÉ (CM) | | | |

**Supplementary Table S2** Optimization of several key parameters of stacks (*M*, *n* and *m*) on a representative subset of 12 individuals. The number of SNPs assembled loci and polymorphic loci is indicated for each combination of parameters. Data were filtered with the populations script from STACKS following the ‘80% rule’ (i.e., keeping only loci shared by at least 80% of samples). The selected combination is *M*5*n*5*m*3.

| Parameters | Total_loci | Removed_loci | Kept_loci | Sites | Filtered_sites | SNPs | Assembled loci | Polymorphic loci | New polymorphic loci |
| --- | --- | --- | --- | --- | --- | --- | --- | --- | --- |
| M1n1m3 | 308193 | 299849 | 8344 | 1454734 | 579 | 8722 | 8344 | 4558 | NA |
| M2n1m3 | 298734 | 290144 | 8590 | 1501638 | 865 | 11679 | 8590 | 5125 | 567 |
| M2n2m3 | 294040 | 285038 | 9002 | 1569683 | 1005 | 13302 | 9002 | 5484 | 359 |
| M3n3m3 | 286787 | 277517 | 9270 | 1619927 | 1367 | 16359 | 9270 | 5871 | 387 |
| M4n4m3 | 282339 | 272975 | 9364 | 1638476 | 1755 | 18254 | 9364 | 6036 | 165 |
| M5n5m3 | 279412 | 269982 | 9430 | 1649788 | 2136 | 19446 | **9430** | 6143 | **107** |
| M6n6m3 | 277235 | 267812 | 9423 | 1650153 | 2396 | 20240 | 9423 | 6170 | 27 |
| M7n7m3 | 275615 | 266236 | 9379 | 1643878 | 2677 | 20774 | 9379 | 6181 | 11 |
| M8n8m3 | 274342 | 264991 | 9351 | 1639061 | 2788 | 21369 | 9351 | **6204** | 23 |
| M9n9m3 | 273351 | 264046 | 9305 | 1631975 | 3061 | 21488 | 9305 | 6172 | -32 |

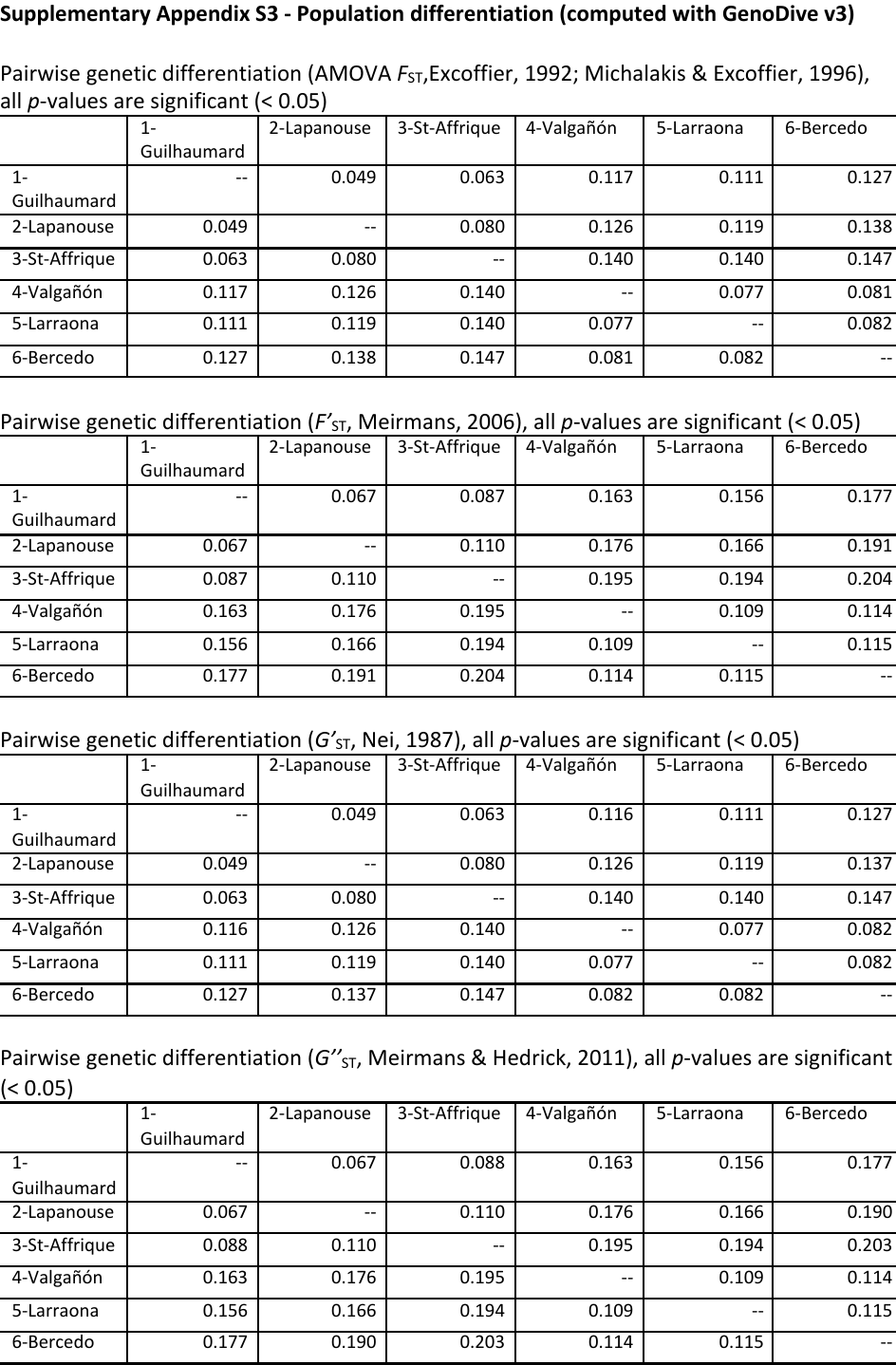

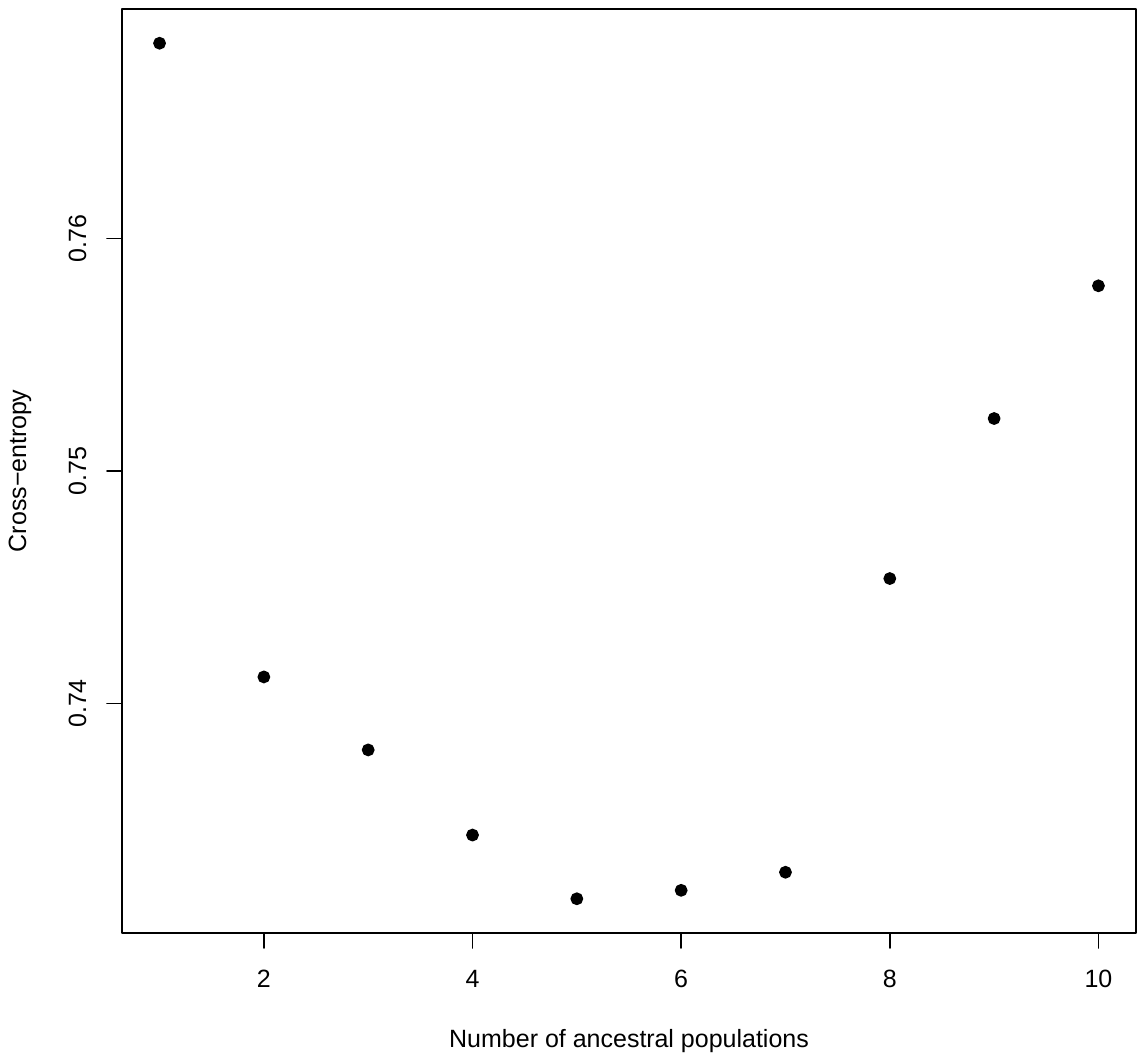

Values of the cross-entropy criterion for a number of clusters ranging from *K* = 1 to 10 (10 sNMF runs each). The optimal number of *K* was found to be 5.

**Supplementary Appendix S4** - **Demographic inference with DILS**

Summary of the different model tested and associated Posterior Probabilities with DILS based on a two-population model. Population considered for the two-populations model, model favored: (ongoing) migration *vs.* (current) isolation (and associated posteriori probability), submodel favored within the (ongoing) migration model: *Isolation Migration* (*IM*) *vs.* *Secondary contact* (*SC*) (and associated probability), model favored between homogeneous and heterogenous introgression rates across the genome (and associated posterior probability), model favored between homogeneous and heterogeneous effective population size across the genome (and associated posterior probability).

| **Pop. 1** | **Pop. 2** | **Migration *vs.* Isolation** | **Post. Prob.** | **IM *vs.* SC** | **Post. Prob.** | ***M*-homo *vs.* *M*-hetero** | **Post. Prob.** | ***N*-homo *vs.* *N*-hetero** | **Post. Prob.** |
| --- | --- | --- | --- | --- | --- | --- | --- | --- | --- |
| *O. aveyronensis* | *O. vitorica* | Migration | 0.591058 | IM | 0.48765 | Mhomo | 0.51008 | Nhetero | 0.86547 |
| Guilhaumard | Valgañón | Migration | 0.57202 | IM | 0.54732 | Mhomo | 0.53402 | Nhetero | 0.82657 |
| Guilhaumard | Larraona | Migration | 0.55163 | IM | 0.53715 | Mhomo | 0.54912 | Nhetero | 0.78942 |
| Guilhaumard | Bercedo | Migration | 0.57202 | IM | 0.54732 | Mhomo | 0.53402 | Nhetero | 0.82657 |
| Lapanouse | Valgañón | Migration | 0.62038 | IM | 0.61995 | Mhetero | 0.53182 | Nhetero | 0.75962 |
| Lapanouse | Larraona | Migration | 0.5732 | IM | 0.4846 | Mhomo | 0.48237 | Nhetero | 0.832 |
| Lapanouse | Bercedo | Migration | 0.66365 | IM | 0.512 | Mhomo | 0.55515 | Nhetero | 0.81382 |
| St-Affrique | Valgañón | Migration | 0.59152 | IM | 0.58022 | Mhomo | 0.49287 | Nhetero | 0.80003 |
| St-Affrique | Larraona | Migration | 0.55713 | IM | 0.52232 | Mhomo | 0.49895 | Nhetero | 0.82562 |
| St-Affrique | Bercedo | Migration | 0.66187 | IM | 0.5744 | Mhomo | 0.53588 | Nhetero | 0.85223 |

^*^In the first raw, all three populations of *O. aveyronesis* subsp. *aveyronensis* (1-Guilhaumard, 2-Lapanouse-de-Cernon, 3-Saint-Affrique) and all three populations of *O. aveyronensis* subsp. *vitorica* (4-Valgañón, 5-Larraona, 6-Bercedo) were grouped by subspecies/country

**Supplementary Appendix S5 – Ecological Niche Modelling with ENMwizard**

Table of the 65 occurrences retrieved from either gbif, iNaturalist (on 2022-07-22) or personal observations and used for Ecological Niche Modelling analyses after spatial thinning (5 km).

| Subspecies | Longitude | Latitude |
| --- | --- | --- |
| *O. a.* subsp. *vitorica* | -2.275645 | 42.7804434 |
| *O. a.* subsp. *vitorica* | -2.40332 | 42.767595 |
| *O. a.* subsp. *vitorica* | -2.45122648561323 | 42.6788794222166 |
| *O. a.* subsp. *vitorica* | -2.5249 | 42.7286 |
| *O. a.* subsp. *vitorica* | -3.02 | 42.330002 |
| *O. a.* subsp. *vitorica* | -3.031 | 42.997 |
| *O. a.* subsp. *vitorica* | -3.08124464522541 | 42.9588790264644 |
| *O. a.* subsp. *vitorica* | -3.0816384 | 42.3188134 |
| *O. a.* subsp. *vitorica* | -3.215 | 43.105 |
| *O. a.* subsp. *vitorica* | -3.239 | 42.889 |
| *O. a.* subsp. *vitorica* | -3.38 | 43.119999 |
| *O. a.* subsp. *vitorica* | -3.4350033 | 43.0943383 |
| *O. a.* subsp. *aveyronensis* | 2.65499 | 44.01772 |
| *O. a.* subsp. *aveyronensis* | 2.67968 | 44.28031 |
| *O. a.* subsp. *aveyronensis* | 2.70838 | 44.46752 |
| *O. a.* subsp. *aveyronensis* | 2.76421 | 44.51908 |
| *O. a.* subsp. *aveyronensis* | 2.811540039 | 43.993294286 |
| *O. a.* subsp. *aveyronensis* | 2.8322141529 | 43.8976433969 |
| *O. a.* subsp. *aveyronensis* | 2.8346950947 | 43.8492039467 |
| *O. a.* subsp. *aveyronensis* | 2.83911 | 44.0857 |
| *O. a.* subsp. *aveyronensis* | 2.8575999891 | 44.1851965579 |
| *O. a.* subsp. *aveyronensis* | 2.86679 | 43.96566 |
| *O. a.* subsp. *aveyronensis* | 2.875407 | 44.029602 |
| *O. a.* subsp. *aveyronensis* | 2.8783124487 | 44.1287150945 |
| *O. a.* subsp. *aveyronensis* | 2.90319 | 44.36188 |
| *O. a.* subsp. *aveyronensis* | 2.9222541615 | 43.8045169412 |
| *O. a.* subsp. *aveyronensis* | 2.9232277654 | 44.0779395852 |
| *O. a.* subsp. *aveyronensis* | 2.9268583 | 43.9807617 |
| *O. a.* subsp. *aveyronensis* | 2.9298327438 | 43.9314120385 |
| *O. a.* subsp. *aveyronensis* | 2.93744 | 44.20399 |
| *O. a.* subsp. *aveyronensis* | 2.9547298317 | 44.1420895921 |
| *O. a.* subsp. *aveyronensis* | 3.0002098476 | 43.8542660825 |
| *O. a.* subsp. *aveyronensis* | 3.0020481478 | 43.9457784661 |
| *O. a.* subsp. *aveyronensis* | 3.003396 | 44.001232 |
| *O. a.* subsp. *aveyronensis* | 3.0345941637 | 43.8048848807 |
| *O. a.* subsp. *aveyronensis* | 3.046845611 | 44.0483102478 |
| *O. a.* subsp. *aveyronensis* | 3.0479867409 | 43.887067742 |
| *O. a.* subsp. *aveyronensis* | 3.05101 | 44.21664 |
| *O. a.* subsp. *aveyronensis* | 3.0826525347 | 44.1453995363 |
| *O. a.* subsp. *aveyronensis* | 3.08389 | 44.30411 |
| *O. a.* subsp. *aveyronensis* | 3.0904516 | 43.9893234 |
| *O. a.* subsp. *aveyronensis* | 3.0912128334 | 43.9278601562 |
| *O. a.* subsp. *aveyronensis* | 3.1183010628 | 43.8762519387 |
| *O. a.* subsp. *aveyronensis* | 3.1361367632 | 43.8024363359 |
| *O. a.* subsp. *aveyronensis* | 3.138695676 | 44.0226115569 |
| *O. a.* subsp. *aveyronensis* | 3.1460843178 | 43.9665359298 |
| *O. a.* subsp. *aveyronensis* | 3.18621 | 43.75386 |
| *O. a.* subsp. *aveyronensis* | 3.1939584 | 43.8459333 |
| *O. a.* subsp. *aveyronensis* | 3.20067 | 43.89242 |
| *O. a.* subsp. *aveyronensis* | 3.20818 | 43.97388 |
| *O. a.* subsp. *aveyronensis* | 3.2414629697 | 44.376710497 |
| *O. a.* subsp. *aveyronensis* | 3.2560921977 | 44.2923954426 |
| *O. a.* subsp. *aveyronensis* | 3.2613017274 | 43.9484299637 |
| *O. a.* subsp. *aveyronensis* | 3.265965 | 43.897305 |
| *O. a.* subsp. *aveyronensis* | 3.2686741302 | 44.2008593735 |
| *O. a.* subsp. *aveyronensis* | 3.2697284783 | 43.8421894317 |
| *O. a.* subsp. *aveyronensis* | 3.27985 | 43.79138 |
| *O. a.* subsp. *aveyronensis* | 3.2799926283 | 43.9945204045 |
| *O. a.* subsp. *aveyronensis* | 3.3337232866 | 43.9680225415 |
| *O. a.* subsp. *aveyronensis* | 3.35181 | 43.77966 |
| *O. a.* subsp. *aveyronensis* | 3.3555863589 | 43.8320425048 |
| *O. a.* subsp. *aveyronensis* | 3.3666625611 | 44.3386667675 |
| *O. a.* subsp. *aveyronensis* | 3.36738 | 43.57971 |
| *O. a.* subsp. *aveyronensis* | 3.3729085136 | 44.2386451921 |
| *O. a.* subsp. *aveyronensis* | 3.3767178499 | 43.8986100969 |

^*^GPS coordinates are given in WGS84.

Summary of the 135 models of Maxent ENMs generated by ENMeval. The best model is indicated in bold.

|  | **Settings** | **feat.** | **rm** | **train.AUC** | **avg.test.AUC** | **var.test.AUC** | **avg.diff.AUC** | **var.diff.AUC** | **avg.test.orMTP** | **var.test.orMTP** | **avg.test.or10pct** | **var.test.or10pct** | **AICc** | **delta.AICc** | **w.AIC** | **Param.** | **sel.cri** | **mod.nms** | **rankAIC** |
| --- | --- | --- | --- | --- | --- | --- | --- | --- | --- | --- | --- | --- | --- | --- | --- | --- | --- | --- | --- |
| 1 | L_0,5 | L | 0,5 | 0,749 | 0,713 | 0,088 | 0,103 | 0,072 | 0,147 | 0,087 | 0,162 | 0,105 | 1000,621 | 75,601 | 0,000 | 7 |  | Mod_0,5_L | 106 |
| 2 | P_0,5 | P | 0,5 | 0,785 | 0,802 | 0,015 | 0,021 | 0,002 | 0,029 | 0,003 | 0,029 | 0,003 | 986,327 | 61,308 | 0,000 | 10 |  | Mod_0,5_P | 76 |
| 3 | Q_0,5 | Q | 0,5 | 0,742 | 0,698 | 0,081 | 0,104 | 0,064 | 0,147 | 0,087 | 0,162 | 0,105 | 1003,903 | 78,884 | 0,000 | 7 |  | Mod_0,5_Q | 133 |
| 4 | H_0,5 | H | 0,5 | 0,901 | 0,821 | 0,060 | 0,094 | 0,061 | 0,133 | 0,051 | 0,225 | 0,062 | 991,668 | 66,649 | 0,000 | 32 |  | Mod_0,5_H | 81 |
| 5 | LP_0,5 | LP | 0,5 | 0,783 | 0,759 | 0,064 | 0,087 | 0,062 | 0,147 | 0,087 | 0,162 | 0,105 | 990,138 | 65,119 | 0,000 | 11 |  | Mod_0,5_LP | 77 |
| 6 | LQ_0,5 | LQ | 0,5 | 0,825 | 0,783 | 0,056 | 0,072 | 0,045 | 0,088 | 0,031 | 0,119 | 0,039 | 968,081 | 43,062 | 0,000 | 11 |  | Mod_0,5_LQ | 67 |
| 7 | LH_0,5 | LH | 0,5 | 0,900 | 0,818 | 0,060 | 0,097 | 0,062 | 0,133 | 0,051 | 0,241 | 0,056 | 991,663 | 66,643 | 0,000 | 32 |  | Mod_0,5_LH | 80 |
| 8 | PQ_0,5 | PQ | 0,5 | 0,810 | 0,789 | 0,052 | 0,074 | 0,049 | 0,103 | 0,042 | 0,163 | 0,081 | 974,257 | 49,237 | 0,000 | 11 |  | Mod_0,5_PQ | 70 |
| 9 | PH_0,5 | PH | 0,5 | 0,901 | 0,819 | 0,061 | 0,098 | 0,062 | 0,133 | 0,051 | 0,225 | 0,062 | 968,840 | 43,820 | 0,000 | 29 |  | Mod_0,5_PH | 68 |
| 10 | QH_0,5 | QH | 0,5 | 0,901 | 0,819 | 0,059 | 0,095 | 0,060 | 0,133 | 0,051 | 0,241 | 0,056 | 1008,166 | 83,146 | 0,000 | 34 |  | Mod_0,5_QH | 134 |
| 11 | LPQ_0,5 | LPQ | 0,5 | 0,811 | 0,784 | 0,059 | 0,079 | 0,056 | 0,103 | 0,042 | 0,163 | 0,081 | 980,293 | 55,273 | 0,000 | 13 |  | Mod_0,5_LPQ | 72 |
| 12 | LPH_0,5 | LPH | 0,5 | 0,901 | 0,819 | 0,061 | 0,097 | 0,062 | 0,133 | 0,051 | 0,225 | 0,062 | 968,840 | 43,820 | 0,000 | 29 |  | Mod_0,5_LPH | 69 |
| 13 | LQH_0,5 | LQH | 0,5 | 0,900 | 0,819 | 0,059 | 0,094 | 0,059 | 0,133 | 0,051 | 0,241 | 0,056 | 1008,908 | 83,888 | 0,000 | 34 |  | Mod_0,5_LQH | 135 |
| 14 | PQH_0,5 | PQH | 0,5 | 0,901 | 0,819 | 0,061 | 0,097 | 0,062 | 0,133 | 0,051 | 0,225 | 0,062 | 956,072 | 31,052 | 0,000 | 27 |  | Mod_0,5_PQH | 65 |
| 15 | LPQH_0,5 | LPQH | 0,5 | 0,901 | 0,819 | 0,061 | 0,097 | 0,062 | 0,133 | 0,051 | 0,225 | 0,062 | 961,878 | 36,858 | 0,000 | 28 |  | Mod_0,5_LPQH | 66 |
| 16 | L_1 | L | 1 | 0,748 | 0,714 | 0,088 | 0,103 | 0,075 | 0,162 | 0,105 | 0,162 | 0,105 | 1000,888 | 75,868 | 0,000 | 7 |  | Mod_1,0_L | 107 |
| 17 | P_1 | P | 1 | 0,772 | 0,775 | 0,018 | 0,020 | 0,002 | 0,029 | 0,003 | 0,029 | 0,003 | 990,656 | 65,636 | 0,000 | 8 |  | Mod_1,0_P | 79 |
| 18 | Q_1 | Q | 1 | 0,741 | 0,699 | 0,080 | 0,102 | 0,068 | 0,147 | 0,087 | 0,162 | 0,105 | 1001,560 | 76,540 | 0,000 | 6 |  | Mod_1,0_Q | 113 |
| 19 | H_1 | H | 1 | 0,890 | 0,832 | 0,055 | 0,084 | 0,052 | 0,104 | 0,028 | 0,149 | 0,049 | 932,515 | 7,495 | 0,005 | 20 |  | Mod_1,0_H | 23 |
| 20 | LP_1 | LP | 1 | 0,769 | 0,744 | 0,063 | 0,092 | 0,061 | 0,147 | 0,087 | 0,162 | 0,105 | 990,635 | 65,616 | 0,000 | 8 |  | Mod_1,0_LP | 78 |
| 21 | LQ_1 | LQ | 1 | 0,804 | 0,735 | 0,086 | 0,099 | 0,070 | 0,147 | 0,087 | 0,162 | 0,105 | 978,534 | 53,515 | 0,000 | 10 |  | Mod_1,0_LQ | 71 |
| 22 | LH_1 | LH | 1 | 0,886 | 0,829 | 0,057 | 0,085 | 0,056 | 0,133 | 0,051 | 0,149 | 0,049 | 930,637 | 5,617 | 0,014 | 19 |  | Mod_1,0_LH | 12 |
| 23 | PQ_1 | PQ | 1 | 0,798 | 0,762 | 0,055 | 0,079 | 0,057 | 0,103 | 0,042 | 0,162 | 0,105 | 982,094 | 57,074 | 0,000 | 10 |  | Mod_1,0_PQ | 74 |
| 24 | PH_1 | PH | 1 | 0,888 | 0,834 | 0,050 | 0,081 | 0,044 | 0,104 | 0,028 | 0,165 | 0,049 | 925,128 | 0,109 | 0,218 | 18 |  | Mod_1,0_PH | 2 |
| 25 | QH_1 | QH | 1 | 0,886 | 0,830 | 0,058 | 0,084 | 0,054 | 0,133 | 0,051 | 0,149 | 0,049 | 930,826 | 5,806 | 0,013 | 19 |  | Mod_1,0_QH | 13 |
| 26 | LPQ_1 | LPQ | 1 | 0,798 | 0,757 | 0,061 | 0,085 | 0,065 | 0,103 | 0,042 | 0,162 | 0,105 | 982,039 | 57,019 | 0,000 | 10 |  | Mod_1,0_LPQ | 73 |
| 27 | LPH_1 | LPH | 1 | 0,888 | 0,834 | 0,049 | 0,081 | 0,043 | 0,104 | 0,028 | 0,165 | 0,049 | 928,965 | 3,945 | 0,032 | 19 |  | Mod_1,0_LPH | 8 |
| 28 | LQH_1 | LQH | 1 | 0,885 | 0,829 | 0,057 | 0,084 | 0,054 | 0,133 | 0,051 | 0,149 | 0,049 | 931,047 | 6,028 | 0,011 | 19 |  | Mod_1,0_LQH | 15 |
| **29** | **PQH_1** | **PQH** | **1** | **0,888** | **0,834** | **0,050** | **0,081** | **0,044** | **0,104** | **0,028** | **0,165** | **0,049** | **925,020** | **0,000** | **0,230** | **18** | **LowAIC** | **Mod_1,0_PQH** | **1** |
| 30 | LPQH_1 | LPQH | 1 | 0,889 | 0,834 | 0,050 | 0,081 | 0,044 | 0,104 | 0,028 | 0,165 | 0,049 | 937,221 | 12,201 | 0,001 | 21 |  | Mod_1,0_LPQH | 36 |
| 31 | L_1,5 | L | 1,5 | 0,748 | 0,715 | 0,088 | 0,104 | 0,079 | 0,162 | 0,105 | 0,162 | 0,105 | 998,618 | 73,599 | 0,000 | 6 |  | Mod_1,5_L | 91 |
| 32 | P_1,5 | P | 1,5 | 0,750 | 0,761 | 0,020 | 0,015 | 0,002 | 0,029 | 0,003 | 0,029 | 0,003 | 1001,809 | 76,790 | 0,000 | 8 |  | Mod_1,5_P | 117 |
| 33 | Q_1,5 | Q | 1,5 | 0,741 | 0,702 | 0,077 | 0,099 | 0,071 | 0,147 | 0,087 | 0,162 | 0,105 | 1001,802 | 76,783 | 0,000 | 6 |  | Mod_1,5_Q | 116 |
| 34 | H_1,5 | H | 1,5 | 0,881 | 0,846 | 0,038 | 0,067 | 0,033 | 0,074 | 0,012 | 0,149 | 0,049 | 938,791 | 13,771 | 0,000 | 19 |  | Mod_1,5_H | 40 |
| 35 | LP_1,5 | LP | 1,5 | 0,754 | 0,738 | 0,063 | 0,092 | 0,062 | 0,147 | 0,087 | 0,162 | 0,105 | 996,912 | 71,892 | 0,000 | 7 |  | Mod_1,5_LP | 85 |
| 36 | LQ_1,5 | LQ | 1,5 | 0,784 | 0,723 | 0,082 | 0,096 | 0,072 | 0,147 | 0,087 | 0,162 | 0,105 | 985,475 | 60,455 | 0,000 | 8 |  | Mod_1,5_LQ | 75 |
| 37 | LH_1,5 | LH | 1,5 | 0,877 | 0,841 | 0,042 | 0,066 | 0,036 | 0,104 | 0,028 | 0,133 | 0,051 | 925,646 | 0,627 | 0,168 | 15 |  | Mod_1,5_LH | 3 |
| 38 | PQ_1,5 | PQ | 1,5 | 0,780 | 0,746 | 0,056 | 0,082 | 0,059 | 0,132 | 0,070 | 0,162 | 0,105 | 992,900 | 67,880 | 0,000 | 10 |  | Mod_1,5_PQ | 84 |
| 39 | PH_1,5 | PH | 1,5 | 0,878 | 0,852 | 0,030 | 0,049 | 0,016 | 0,045 | 0,003 | 0,134 | 0,038 | 928,683 | 3,664 | 0,037 | 16 |  | Mod_1,5_PH | 6 |
| 40 | QH_1,5 | QH | 1,5 | 0,877 | 0,842 | 0,043 | 0,063 | 0,033 | 0,104 | 0,028 | 0,133 | 0,051 | 937,011 | 11,992 | 0,001 | 18 |  | Mod_1,5_QH | 29 |
| 41 | LPQ_1,5 | LPQ | 1,5 | 0,780 | 0,741 | 0,062 | 0,087 | 0,066 | 0,132 | 0,070 | 0,162 | 0,105 | 992,770 | 67,750 | 0,000 | 10 |  | Mod_1,5_LPQ | 83 |
| 42 | LPH_1,5 | LPH | 1,5 | 0,878 | 0,851 | 0,030 | 0,049 | 0,015 | 0,045 | 0,003 | 0,134 | 0,038 | 928,699 | 3,680 | 0,037 | 16 |  | Mod_1,5_LPH | 7 |
| 43 | LQH_1,5 | LQH | 1,5 | 0,877 | 0,845 | 0,038 | 0,058 | 0,028 | 0,104 | 0,028 | 0,133 | 0,051 | 929,625 | 4,605 | 0,023 | 16 |  | Mod_1,5_LQH | 9 |
| 44 | PQH_1,5 | PQH | 1,5 | 0,878 | 0,851 | 0,030 | 0,049 | 0,015 | 0,045 | 0,003 | 0,134 | 0,038 | 928,658 | 3,638 | 0,037 | 16 |  | Mod_1,5_PQH | 4 |
| 45 | LPQH_1,5 | LPQH | 1,5 | 0,878 | 0,852 | 0,030 | 0,049 | 0,015 | 0,045 | 0,003 | 0,134 | 0,038 | 928,658 | 3,638 | 0,037 | 16 |  | Mod_1,5_LPQH | 5 |
| 46 | L_2 | L | 2 | 0,747 | 0,716 | 0,087 | 0,104 | 0,083 | 0,162 | 0,105 | 0,162 | 0,105 | 998,853 | 73,834 | 0,000 | 6 |  | Mod_2,0_L | 93 |
| 47 | P_2 | P | 2 | 0,743 | 0,755 | 0,022 | 0,012 | 0,001 | 0,029 | 0,003 | 0,029 | 0,003 | 1002,681 | 77,661 | 0,000 | 6 |  | Mod_2,0_P | 126 |
| 48 | Q_2 | Q | 2 | 0,739 | 0,706 | 0,075 | 0,095 | 0,073 | 0,147 | 0,087 | 0,162 | 0,105 | 1002,136 | 77,117 | 0,000 | 6 |  | Mod_2,0_Q | 122 |
| 49 | H_2 | H | 2 | 0,872 | 0,859 | 0,025 | 0,049 | 0,018 | 0,060 | 0,007 | 0,133 | 0,051 | 930,512 | 5,493 | 0,015 | 14 |  | Mod_2,0_H | 11 |
| 50 | LP_2 | LP | 2 | 0,751 | 0,740 | 0,056 | 0,085 | 0,056 | 0,118 | 0,055 | 0,147 | 0,087 | 1001,125 | 76,106 | 0,000 | 8 |  | Mod_2,0_LP | 111 |
| 51 | LQ_2 | LQ | 2 | 0,764 | 0,718 | 0,078 | 0,095 | 0,076 | 0,147 | 0,087 | 0,162 | 0,105 | 992,528 | 67,508 | 0,000 | 7 |  | Mod_2,0_LQ | 82 |
| 52 | LH_2 | LH | 2 | 0,865 | 0,849 | 0,031 | 0,047 | 0,020 | 0,045 | 0,003 | 0,119 | 0,039 | 929,952 | 4,933 | 0,020 | 13 |  | Mod_2,0_LH | 10 |
| 53 | PQ_2 | PQ | 2 | 0,759 | 0,748 | 0,049 | 0,077 | 0,053 | 0,132 | 0,070 | 0,147 | 0,087 | 1000,015 | 74,996 | 0,000 | 9 |  | Mod_2,0_PQ | 102 |
| 54 | PH_2 | PH | 2 | 0,867 | 0,848 | 0,032 | 0,028 | 0,005 | 0,045 | 0,003 | 0,045 | 0,003 | 936,229 | 11,209 | 0,001 | 15 |  | Mod_2,0_PH | 27 |
| 55 | QH_2 | QH | 2 | 0,867 | 0,853 | 0,030 | 0,043 | 0,017 | 0,060 | 0,007 | 0,119 | 0,039 | 939,312 | 14,293 | 0,000 | 16 |  | Mod_2,0_QH | 41 |
| 56 | LPQ_2 | LPQ | 2 | 0,759 | 0,743 | 0,056 | 0,082 | 0,061 | 0,147 | 0,087 | 0,162 | 0,105 | 999,823 | 74,804 | 0,000 | 9 |  | Mod_2,0_LPQ | 99 |
| 57 | LPH_2 | LPH | 2 | 0,867 | 0,848 | 0,032 | 0,028 | 0,005 | 0,045 | 0,003 | 0,045 | 0,003 | 932,874 | 7,854 | 0,005 | 14 |  | Mod_2,0_LPH | 25 |
| 58 | LQH_2 | LQH | 2 | 0,866 | 0,854 | 0,028 | 0,038 | 0,013 | 0,045 | 0,003 | 0,104 | 0,028 | 933,329 | 8,309 | 0,004 | 14 |  | Mod_2,0_LQH | 26 |
| 59 | PQH_2 | PQH | 2 | 0,867 | 0,848 | 0,032 | 0,028 | 0,005 | 0,045 | 0,003 | 0,045 | 0,003 | 936,237 | 11,218 | 0,001 | 15 |  | Mod_2,0_PQH | 28 |
| 60 | LPQH_2 | LPQH | 2 | 0,867 | 0,848 | 0,032 | 0,028 | 0,005 | 0,045 | 0,003 | 0,045 | 0,003 | 932,801 | 7,782 | 0,005 | 14 |  | Mod_2,0_LPQH | 24 |
| 61 | L_2,5 | L | 2,5 | 0,747 | 0,718 | 0,087 | 0,104 | 0,087 | 0,162 | 0,105 | 0,176 | 0,125 | 999,151 | 74,132 | 0,000 | 6 |  | Mod_2,5_L | 94 |
| 62 | P_2,5 | P | 2,5 | 0,741 | 0,754 | 0,022 | 0,008 | 0,001 | 0,029 | 0,003 | 0,029 | 0,003 | 1001,598 | 76,578 | 0,000 | 5 |  | Mod_2,5_P | 114 |
| 63 | Q_2,5 | Q | 2,5 | 0,738 | 0,709 | 0,073 | 0,090 | 0,073 | 0,147 | 0,087 | 0,162 | 0,105 | 1002,560 | 77,540 | 0,000 | 6 |  | Mod_2,5_Q | 125 |
| 64 | H_2,5 | H | 2,5 | 0,864 | 0,872 | 0,016 | 0,027 | 0,006 | 0,030 | 0,001 | 0,074 | 0,012 | 937,180 | 12,161 | 0,001 | 13 |  | Mod_2,5_H | 35 |
| 65 | LP_2,5 | LP | 2,5 | 0,747 | 0,741 | 0,047 | 0,075 | 0,049 | 0,118 | 0,055 | 0,147 | 0,087 | 1003,549 | 78,530 | 0,000 | 8 |  | Mod_2,5_LP | 130 |
| 66 | LQ_2,5 | LQ | 2,5 | 0,750 | 0,721 | 0,075 | 0,092 | 0,076 | 0,147 | 0,087 | 0,162 | 0,105 | 996,981 | 71,961 | 0,000 | 6 |  | Mod_2,5_LQ | 86 |
| 67 | LH_2,5 | LH | 2,5 | 0,856 | 0,846 | 0,034 | 0,047 | 0,020 | 0,045 | 0,003 | 0,119 | 0,039 | 944,638 | 19,618 | 0,000 | 15 |  | Mod_2,5_LH | 53 |
| 68 | PQ_2,5 | PQ | 2,5 | 0,749 | 0,752 | 0,042 | 0,070 | 0,044 | 0,118 | 0,055 | 0,147 | 0,087 | 1001,882 | 76,862 | 0,000 | 8 |  | Mod_2,5_PQ | 119 |
| 69 | PH_2,5 | PH | 2,5 | 0,859 | 0,849 | 0,034 | 0,023 | 0,004 | 0,045 | 0,003 | 0,045 | 0,003 | 931,047 | 6,027 | 0,011 | 11 |  | Mod_2,5_PH | 14 |
| 70 | QH_2,5 | QH | 2,5 | 0,857 | 0,850 | 0,033 | 0,045 | 0,018 | 0,060 | 0,007 | 0,119 | 0,039 | 945,510 | 20,491 | 0,000 | 15 |  | Mod_2,5_QH | 60 |
| 71 | LPQ_2,5 | LPQ | 2,5 | 0,749 | 0,747 | 0,047 | 0,077 | 0,053 | 0,118 | 0,055 | 0,147 | 0,087 | 1001,901 | 76,881 | 0,000 | 8 |  | Mod_2,5_LPQ | 120 |
| 72 | LPH_2,5 | LPH | 2,5 | 0,859 | 0,849 | 0,034 | 0,023 | 0,004 | 0,045 | 0,003 | 0,045 | 0,003 | 931,049 | 6,030 | 0,011 | 11 |  | Mod_2,5_LPH | 16 |
| 73 | LQH_2,5 | LQH | 2,5 | 0,855 | 0,854 | 0,028 | 0,037 | 0,012 | 0,045 | 0,003 | 0,104 | 0,028 | 942,112 | 17,092 | 0,000 | 14 |  | Mod_2,5_LQH | 50 |
| 74 | PQH_2,5 | PQH | 2,5 | 0,859 | 0,849 | 0,034 | 0,023 | 0,004 | 0,045 | 0,003 | 0,045 | 0,003 | 931,102 | 6,082 | 0,011 | 11 |  | Mod_2,5_PQH | 18 |
| 75 | LPQH_2,5 | LPQH | 2,5 | 0,859 | 0,849 | 0,034 | 0,023 | 0,004 | 0,045 | 0,003 | 0,045 | 0,003 | 931,064 | 6,044 | 0,011 | 11 |  | Mod_2,5_LPQH | 17 |
| 76 | L_3 | L | 3 | 0,746 | 0,720 | 0,085 | 0,101 | 0,086 | 0,162 | 0,105 | 0,176 | 0,125 | 999,508 | 74,488 | 0,000 | 6 |  | Mod_3,0_L | 95 |
| 77 | P_3 | P | 3 | 0,740 | 0,753 | 0,022 | 0,003 | 0,000 | 0,029 | 0,003 | 0,029 | 0,003 | 1002,238 | 77,218 | 0,000 | 5 |  | Mod_3,0_P | 124 |
| 78 | Q_3 | Q | 3 | 0,736 | 0,711 | 0,071 | 0,088 | 0,070 | 0,147 | 0,087 | 0,162 | 0,105 | 1003,072 | 78,052 | 0,000 | 6 |  | Mod_3,0_Q | 129 |
| 79 | H_3 | H | 3 | 0,859 | 0,872 | 0,015 | 0,021 | 0,004 | 0,030 | 0,001 | 0,074 | 0,012 | 938,749 | 13,730 | 0,000 | 11 |  | Mod_3,0_H | 39 |
| 80 | LP_3 | LP | 3 | 0,744 | 0,746 | 0,035 | 0,061 | 0,033 | 0,088 | 0,031 | 0,118 | 0,055 | 1000,014 | 74,995 | 0,000 | 6 |  | Mod_3,0_LP | 101 |
| 81 | LQ_3 | LQ | 3 | 0,749 | 0,722 | 0,074 | 0,090 | 0,073 | 0,147 | 0,087 | 0,162 | 0,105 | 997,487 | 72,468 | 0,000 | 6 |  | Mod_3,0_LQ | 87 |
| 82 | LH_3 | LH | 3 | 0,846 | 0,846 | 0,037 | 0,049 | 0,021 | 0,045 | 0,003 | 0,119 | 0,039 | 945,929 | 20,909 | 0,000 | 13 |  | Mod_3,0_LH | 61 |
| 83 | PQ_3 | PQ | 3 | 0,746 | 0,757 | 0,032 | 0,056 | 0,028 | 0,074 | 0,022 | 0,118 | 0,055 | 1001,081 | 76,061 | 0,000 | 7 |  | Mod_3,0_PQ | 110 |
| 84 | PH_3 | PH | 3 | 0,853 | 0,850 | 0,035 | 0,021 | 0,004 | 0,045 | 0,003 | 0,045 | 0,003 | 931,504 | 6,485 | 0,009 | 9 |  | Mod_3,0_PH | 19 |
| 85 | QH_3 | QH | 3 | 0,847 | 0,850 | 0,035 | 0,045 | 0,018 | 0,060 | 0,007 | 0,119 | 0,039 | 938,585 | 13,565 | 0,000 | 10 |  | Mod_3,0_QH | 38 |
| 86 | LPQ_3 | LPQ | 3 | 0,746 | 0,749 | 0,040 | 0,069 | 0,042 | 0,118 | 0,055 | 0,147 | 0,087 | 1001,080 | 76,061 | 0,000 | 7 |  | Mod_3,0_LPQ | 109 |
| 87 | LPH_3 | LPH | 3 | 0,853 | 0,850 | 0,035 | 0,021 | 0,004 | 0,045 | 0,003 | 0,045 | 0,003 | 931,504 | 6,485 | 0,009 | 9 |  | Mod_3,0_LPH | 20 |
| 88 | LQH_3 | LQH | 3 | 0,845 | 0,854 | 0,030 | 0,036 | 0,012 | 0,045 | 0,003 | 0,104 | 0,028 | 947,511 | 22,491 | 0,000 | 13 |  | Mod_3,0_LQH | 62 |
| 89 | PQH_3 | PQH | 3 | 0,853 | 0,850 | 0,035 | 0,020 | 0,004 | 0,045 | 0,003 | 0,045 | 0,003 | 931,530 | 6,510 | 0,009 | 9 |  | Mod_3,0_PQH | 21 |
| 90 | LPQH_3 | LPQH | 3 | 0,853 | 0,850 | 0,035 | 0,020 | 0,004 | 0,045 | 0,003 | 0,045 | 0,003 | 931,530 | 6,510 | 0,009 | 9 |  | Mod_3,0_LPQH | 22 |
| 91 | L_3,5 | L | 3,5 | 0,745 | 0,721 | 0,083 | 0,098 | 0,085 | 0,162 | 0,105 | 0,176 | 0,125 | 999,924 | 74,904 | 0,000 | 6 |  | Mod_3,5_L | 100 |
| 92 | P_3,5 | P | 3,5 | 0,739 | 0,753 | 0,023 | 0,006 | 0,000 | 0,029 | 0,003 | 0,029 | 0,003 | 1002,968 | 77,949 | 0,000 | 5 |  | Mod_3,5_P | 128 |
| 93 | Q_3,5 | Q | 3,5 | 0,734 | 0,713 | 0,069 | 0,087 | 0,068 | 0,147 | 0,087 | 0,162 | 0,105 | 1003,671 | 78,651 | 0,000 | 6 |  | Mod_3,5_Q | 131 |
| 94 | H_3,5 | H | 3,5 | 0,856 | 0,871 | 0,015 | 0,015 | 0,002 | 0,016 | 0,001 | 0,074 | 0,012 | 939,422 | 14,403 | 0,000 | 9 |  | Mod_3,5_H | 42 |
| 95 | LP_3,5 | LP | 3,5 | 0,742 | 0,752 | 0,028 | 0,042 | 0,016 | 0,029 | 0,003 | 0,074 | 0,022 | 1001,274 | 76,254 | 0,000 | 6 |  | Mod_3,5_LP | 112 |
| 96 | LQ_3,5 | LQ | 3,5 | 0,747 | 0,725 | 0,072 | 0,089 | 0,071 | 0,147 | 0,087 | 0,162 | 0,105 | 998,093 | 73,074 | 0,000 | 6 |  | Mod_3,5_LQ | 88 |
| 97 | LH_3,5 | LH | 3,5 | 0,838 | 0,845 | 0,038 | 0,048 | 0,021 | 0,045 | 0,003 | 0,119 | 0,039 | 944,189 | 19,169 | 0,000 | 10 |  | Mod_3,5_LH | 52 |
| 98 | PQ_3,5 | PQ | 3,5 | 0,743 | 0,756 | 0,028 | 0,036 | 0,012 | 0,029 | 0,003 | 0,059 | 0,014 | 1000,044 | 75,024 | 0,000 | 6 |  | Mod_3,5_PQ | 104 |
| 99 | PH_3,5 | PH | 3,5 | 0,849 | 0,849 | 0,037 | 0,020 | 0,004 | 0,045 | 0,003 | 0,045 | 0,003 | 937,092 | 12,072 | 0,001 | 9 |  | Mod_3,5_PH | 33 |
| 100 | QH_3,5 | QH | 3,5 | 0,841 | 0,850 | 0,035 | 0,044 | 0,017 | 0,045 | 0,003 | 0,119 | 0,039 | 938,534 | 13,514 | 0,000 | 8 |  | Mod_3,5_QH | 37 |
| 101 | LPQ_3,5 | LPQ | 3,5 | 0,743 | 0,754 | 0,030 | 0,047 | 0,020 | 0,044 | 0,008 | 0,088 | 0,031 | 1000,022 | 75,002 | 0,000 | 6 |  | Mod_3,5_LPQ | 103 |
| 102 | LPH_3,5 | LPH | 3,5 | 0,849 | 0,851 | 0,037 | 0,020 | 0,004 | 0,045 | 0,003 | 0,045 | 0,003 | 937,038 | 12,019 | 0,001 | 9 |  | Mod_3,5_LPH | 30 |
| 103 | LQH_3,5 | LQH | 3,5 | 0,839 | 0,856 | 0,028 | 0,032 | 0,009 | 0,045 | 0,003 | 0,089 | 0,020 | 941,652 | 16,632 | 0,000 | 9 |  | Mod_3,5_LQH | 47 |
| 104 | PQH_3,5 | PQH | 3,5 | 0,849 | 0,850 | 0,036 | 0,020 | 0,003 | 0,045 | 0,003 | 0,045 | 0,003 | 937,098 | 12,079 | 0,001 | 9 |  | Mod_3,5_PQH | 34 |
| 105 | LPQH_3,5 | LPQH | 3,5 | 0,849 | 0,851 | 0,036 | 0,020 | 0,003 | 0,045 | 0,003 | 0,045 | 0,003 | 937,070 | 12,050 | 0,001 | 9 |  | Mod_3,5_LPQH | 32 |
| 106 | L_4 | L | 4 | 0,744 | 0,722 | 0,080 | 0,096 | 0,084 | 0,162 | 0,105 | 0,176 | 0,125 | 1000,398 | 75,378 | 0,000 | 6 |  | Mod_4,0_L | 105 |
| 107 | P_4 | P | 4 | 0,737 | 0,753 | 0,025 | 0,012 | 0,001 | 0,029 | 0,003 | 0,029 | 0,003 | 1003,781 | 78,761 | 0,000 | 5 |  | Mod_4,0_P | 132 |
| 108 | Q_4 | Q | 4 | 0,734 | 0,714 | 0,067 | 0,085 | 0,065 | 0,147 | 0,087 | 0,177 | 0,099 | 1001,696 | 76,676 | 0,000 | 5 |  | Mod_4,0_Q | 115 |
| 109 | H_4 | H | 4 | 0,853 | 0,869 | 0,015 | 0,006 | 0,000 | 0,016 | 0,001 | 0,074 | 0,012 | 944,724 | 19,704 | 0,000 | 9 |  | Mod_4,0_H | 54 |
| 110 | LP_4 | LP | 4 | 0,740 | 0,754 | 0,027 | 0,019 | 0,003 | 0,029 | 0,003 | 0,029 | 0,003 | 1002,699 | 77,679 | 0,000 | 6 |  | Mod_4,0_LP | 127 |
| 111 | LQ_4 | LQ | 4 | 0,746 | 0,727 | 0,069 | 0,086 | 0,067 | 0,147 | 0,087 | 0,162 | 0,105 | 998,768 | 73,748 | 0,000 | 6 |  | Mod_4,0_LQ | 92 |
| 112 | LH_4 | LH | 4 | 0,836 | 0,843 | 0,041 | 0,051 | 0,024 | 0,045 | 0,003 | 0,133 | 0,051 | 942,109 | 17,090 | 0,000 | 8 |  | Mod_4,0_LH | 49 |
| 113 | PQ_4 | PQ | 4 | 0,742 | 0,757 | 0,027 | 0,014 | 0,002 | 0,029 | 0,003 | 0,029 | 0,003 | 998,572 | 73,552 | 0,000 | 5 |  | Mod_4,0_PQ | 90 |
| 114 | PH_4 | PH | 4 | 0,845 | 0,849 | 0,038 | 0,021 | 0,004 | 0,045 | 0,003 | 0,045 | 0,003 | 939,664 | 14,644 | 0,000 | 8 |  | Mod_4,0_PH | 45 |
| 115 | QH_4 | QH | 4 | 0,838 | 0,850 | 0,035 | 0,042 | 0,016 | 0,045 | 0,003 | 0,119 | 0,039 | 939,469 | 14,450 | 0,000 | 7 |  | Mod_4,0_QH | 43 |
| 116 | LPQ_4 | LPQ | 4 | 0,742 | 0,754 | 0,029 | 0,027 | 0,007 | 0,029 | 0,003 | 0,029 | 0,003 | 998,561 | 73,541 | 0,000 | 5 |  | Mod_4,0_LPQ | 89 |
| 117 | LPH_4 | LPH | 4 | 0,845 | 0,850 | 0,038 | 0,021 | 0,004 | 0,045 | 0,003 | 0,045 | 0,003 | 939,664 | 14,644 | 0,000 | 8 |  | Mod_4,0_LPH | 46 |
| 118 | LQH_4 | LQH | 4 | 0,837 | 0,855 | 0,029 | 0,033 | 0,010 | 0,045 | 0,003 | 0,104 | 0,028 | 941,858 | 16,838 | 0,000 | 8 |  | Mod_4,0_LQH | 48 |
| 119 | PQH_4 | PQH | 4 | 0,845 | 0,849 | 0,037 | 0,020 | 0,004 | 0,045 | 0,003 | 0,045 | 0,003 | 937,040 | 12,020 | 0,001 | 7 |  | Mod_4,0_PQH | 31 |
| 120 | LPQH_4 | LPQH | 4 | 0,845 | 0,851 | 0,037 | 0,020 | 0,004 | 0,045 | 0,003 | 0,045 | 0,003 | 939,641 | 14,622 | 0,000 | 8 |  | Mod_4,0_LPQH | 44 |
| 121 | L_4,5 | L | 4,5 | 0,742 | 0,722 | 0,077 | 0,095 | 0,081 | 0,162 | 0,105 | 0,162 | 0,105 | 1000,928 | 75,908 | 0,000 | 6 |  | Mod_4,5_L | 108 |
| 122 | P_4,5 | P | 4,5 | 0,737 | 0,752 | 0,027 | 0,016 | 0,002 | 0,029 | 0,003 | 0,029 | 0,003 | 1002,079 | 77,059 | 0,000 | 4 |  | Mod_4,5_P | 121 |
| 123 | Q_4,5 | Q | 4,5 | 0,734 | 0,716 | 0,066 | 0,084 | 0,064 | 0,147 | 0,087 | 0,177 | 0,099 | 1002,161 | 77,141 | 0,000 | 5 |  | Mod_4,5_Q | 123 |
| 124 | H_4,5 | H | 4,5 | 0,851 | 0,868 | 0,014 | 0,000 | 0,000 | 0,016 | 0,001 | 0,045 | 0,003 | 953,444 | 28,424 | 0,000 | 10 |  | Mod_4,5_H | 64 |
| 125 | LP_4,5 | LP | 4,5 | 0,738 | 0,752 | 0,027 | 0,016 | 0,002 | 0,029 | 0,003 | 0,029 | 0,003 | 1001,822 | 76,802 | 0,000 | 5 |  | Mod_4,5_LP | 118 |
| 126 | LQ_4,5 | LQ | 4,5 | 0,744 | 0,728 | 0,067 | 0,084 | 0,064 | 0,147 | 0,087 | 0,162 | 0,105 | 999,529 | 74,509 | 0,000 | 6 |  | Mod_4,5_LQ | 97 |
| 127 | LH_4,5 | LH | 4,5 | 0,834 | 0,839 | 0,043 | 0,053 | 0,026 | 0,045 | 0,003 | 0,133 | 0,051 | 948,086 | 23,066 | 0,000 | 9 |  | Mod_4,5_LH | 63 |
| 128 | PQ_4,5 | PQ | 4,5 | 0,742 | 0,755 | 0,028 | 0,016 | 0,002 | 0,029 | 0,003 | 0,045 | 0,003 | 999,536 | 74,517 | 0,000 | 5 |  | Mod_4,5_PQ | 98 |
| 129 | PH_4,5 | PH | 4,5 | 0,841 | 0,849 | 0,039 | 0,021 | 0,004 | 0,045 | 0,003 | 0,045 | 0,003 | 945,008 | 19,988 | 0,000 | 8 |  | Mod_4,5_PH | 57 |
| 130 | QH_4,5 | QH | 4,5 | 0,837 | 0,848 | 0,035 | 0,043 | 0,016 | 0,045 | 0,003 | 0,119 | 0,039 | 942,488 | 17,468 | 0,000 | 7 |  | Mod_4,5_QH | 51 |
| 131 | LPQ_4,5 | LPQ | 4,5 | 0,742 | 0,755 | 0,028 | 0,016 | 0,002 | 0,029 | 0,003 | 0,045 | 0,003 | 999,525 | 74,505 | 0,000 | 5 |  | Mod_4,5_LPQ | 96 |
| 132 | LPH_4,5 | LPH | 4,5 | 0,841 | 0,850 | 0,039 | 0,021 | 0,004 | 0,045 | 0,003 | 0,045 | 0,003 | 945,021 | 20,001 | 0,000 | 8 |  | Mod_4,5_LPH | 58 |
| 133 | LQH_4,5 | LQH | 4,5 | 0,835 | 0,852 | 0,031 | 0,037 | 0,012 | 0,045 | 0,003 | 0,119 | 0,039 | 945,178 | 20,159 | 0,000 | 8 |  | Mod_4,5_LQH | 59 |
| 134 | PQH_4,5 | PQH | 4,5 | 0,841 | 0,848 | 0,039 | 0,022 | 0,004 | 0,045 | 0,003 | 0,045 | 0,003 | 944,905 | 19,885 | 0,000 | 8 |  | Mod_4,5_PQH | 56 |
| 135 | LPQH_4,5 | LPQH | 4,5 | 0,841 | 0,850 | 0,039 | 0,022 | 0,004 | 0,045 | 0,003 | 0,045 | 0,003 | 944,903 | 19,883 | 0,000 | 8 |  | Mod_4,5_LPQH | 55 |

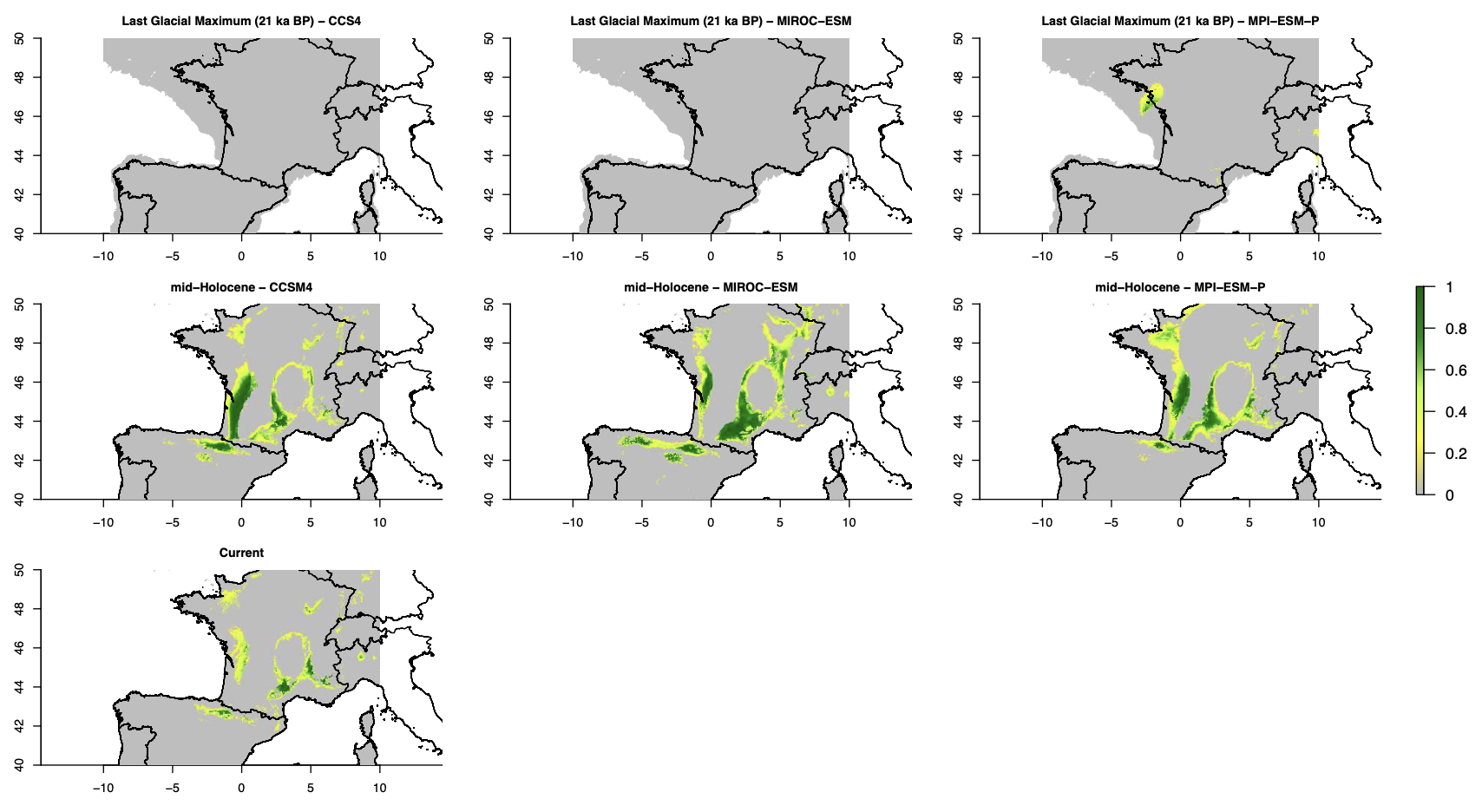

Spatial projections of the bioclimatic niches inferred for the *Ophrys aveyronensis* species complex depicting climate suitability at different time periods from the Last Glacial Maximum (LGM, about 21 kyrs ago) with a different interpolation algorithm (WorldClim instead of CHELSA) and with projections for the mid-Holocene based on different Global Circulation Models (CCSM4, MIROC-ESM and MPI-ESM-P) and currently.
